# Structural and Functional Relevance of Charge Based Transient Interactions inside Intrinsically Disordered Proteins

**DOI:** 10.1101/2024.10.30.621161

**Authors:** Samuel Wohl, Yishai Gilron, Wenwei Zheng

## Abstract

Intrinsically disordered proteins (IDPs) perform a wide range of biological functions without adopting stable, well-defined, three-dimensional structures. Instead, IDPs exist as dynamic ensembles of flexible conformations, traditionally thought to be governed by weak, nonspecific interactions, which are well described by homopolymer theory. However, recent research highlights the presence of transient, specific interactions in several IDPs, suggesting that factors beyond overall size influence their conformational behavior. In this study, we investigate how the spatial arrangement of charged amino acids within IDP sequences shapes the prevalence of transient, specific interactions. Through a series of model peptides, we establish a quantitative empirical relationship between the fraction of transient interactions and a novel sequence metric, termed effective charged patch length, which characterizes the ability of charged patches to drive these interactions. By examining IDP ensembles with varying levels of transient interactions, we further explore their heteropolymeric structural behavior in phase-separated condensates, where we observe the formation of a condensate-spanning network structure. Additionally, we perform a proteome-wide scan for charge-based transient interactions within disordered regions of the human proteome, revealing that approximately 10% of these regions exhibit such charge-driven transient interactions, leading to heteropolymeric behaviors in their conformational ensembles. Finally, we examine how these charge-based transient interactions correlate with molecular functions, identifying specific biological roles in which these interactions are enriched.

---

Intrinsically disordered proteins (IDPs) are involved in a wide array of cellular processes.^1,2^ These include transcription and signaling transduction,^3^ formation of membraneless organelles,^4–6^ and functional or pathological aggregates.^7,8^ However, their lack of a welldefined three dimensional structure and flexible conformations pose significant challenges in their biophysical characterization.^9,10^ IDPs challenge the traditional sequence-structure-function paradigm typically applied to folded proteins.^11,12^ In those cases, structure often serves as an important intermediate, simplifying the high-dimensional sequence space into a more manageable low-dimensional structural representation,^13^ which facilitates the exploration of functional relationships. Instead, IDP sequences give rise to an ensemble of conformations which still contains many degrees of freedom. ^14–18^ This means it is often unclear exactly which ensemble properties are functionally relevant. One major challenge for biophysicists is therefore to look for key structural properties that capture these conformational ensembles,^19–23^ decode the sequence grammar of these structural properties,^24–26^ and finally address how these features contribute to IDP functions. ^27,28^

Thanks to advancements in a variety of experimental technologies, a wealth of structural properties of IDPs can now be directly measured.^29^ These properties can be classified into global conformational, site-specific, or pairwise amino-acid properties. Global conformational properties include radius of gyration from small-angle X-ray scattering (SAXS), ^30^ hydrodynamic radius from dynamic light scattering (DLS) ^31^ pulsed-field gradient nuclear magnetic resonance (PFG-NMR), ^32^ and end-to-end distance from Förster resonance energy transfer (FRET).^33^ These global conformational properties are useful for probing overall structural changes in response to varying environmental conditions, such as temperature, ionic strength, denaturants, and crowding.^34–37^ Consequently, they are often integrated with physics-based amino acid properties in computational models to decode sequence grammars.^23,38,39^ Site-specific properties rely on labeling or assignments, and various nuclear magnetic resonance (NMR) methods are often used. For example, secondary structure chemical shifts provide site-specific secondary structure preferences,^40^ while hydrogen exchange,^41^ hydroxyl radical protein footprinting,^42^ solvent paramagnetic relaxation enhancement (sPRE),^43^ and NMR relaxation parameters^44^ offer insights into the solvent exposure of specific amino acids. Additionally, chemical shift perturbation, when coupled with mutations and/or varying environmental conditions, is a powerful tool for understanding site-specific conformational changes.^45^ Pairwise amino acid properties are assessed by introducing pairs of site-specific labels. These include metrics such as distances from FRET^33^ or paramagnetic relaxation enhancement (PRE),^46^ and relaxation dynamics from nanosecond fluorescence correlation spectroscopy (nsFCS)^47^ or photoinduced electron transfer (PET).^48^

Each of these experimental methods often excel at probing one type of structural property, which might not always be functionally significant. In practice, one can only formulate a hypothesis about the most functionally relevant structure property from within the available space of experimental observables. For instance, one of the most common experimental observables across various IDP sequences is the overall size of an IDP, quantified by either the radius of gyration or the hydrodynamic radius. ^49^ The polymer scaling exponent, based on homopolymer theory, which adjusts the radius of gyration and hydrodynamic radius in relation to protein chain length, is a valuable structural property for characterizing various IDPs.^50,51^ This scaling exponent has been successful in interpreting a range of experimental measurements^52,53^ and in explaining phenomena such as liquid-liquid phase separation (LLPS) of IDPs.^54^ The underlying assumption for using this scaling exponent is that IDPs behave like homopolymers and are dominated by nonspecific, weak interactions. This assumption is accurate in some cases because IDPs often have low complexity sequences, that is a small number of amino acids constitute a large fraction of the sequence and/or the sequence contains repeated fragments. In these cases, conformational and functional behaviors can be explained using only the overall sequence properties. The fraction of aromatic amino acids, for example, has been shown to be important for predicting LLPS behavior.^55^

However, emerging evidence indicates that IDPs are not solely governed by nonspecific, weak interactions, and that their overall size alone is insufficient to fully capture their conformational behavior. In fact, specific interactions between pairs of amino acids or fragments with long sequence separations are often observed in PRE measurements.^56^ Such behavior has also been reported by FRET^36^ and PET,^57^ with multiple pair labeling positions used to check deviations from a homopolymer model. The challenge in recognizing these interactions arises from their transient nature. They may exist only in a small fraction of conformations within the ensemble and/or have short lifetimes. These interactions can be caused by either charged^36,57^ or hydrophobic amino acids. ^58,59^ They can also be either intramolecular, occurring between different regions or segments within the same protein,^28,60^ or intermolecular, occurring between multiple proteins.^61–63^ The strength of these transient interactions is expected to be highly sensitive to environmental conditions or post-translational modifications,^28^ providing a flexible means of adjusting interactions according to the environment. Their presence in various transcriptional activation domains suggests a functional role in transcriptional regulation,^28,60,61,64^ for example, by regulating the binding of the transcriptional activation domain with the coactivator or adjusting the solvent exposure of the DNA binding domain. Furthermore, transient interactions may influence the microstructures within IDP condensates, potentially providing fine-tuning for the formation of membraneless organelles.^65,66^

Site-specific labeling experimental measurements can directly probe these transient, specific interactions, but these come with two primary concerns. First, the label itself might perturb the interactions, as transient interactions are often sensitive to variations in the chemical environment. Second, site-specific labeling requires good empirical knowledge of the interaction sites and often requires the introduction of multiple labeling positions. Computational methods can assist in both aspects: first by testing the impact of a given label^67^ and second by generating an ensemble that best matches experimental data, even in the absence of direct measurements between sites with transient interactions.^14–16,18,59^ However, due to the many degrees of freedom involved, generating an ensemble that matches limited experimental data can involve significant uncertainties. The choice of the physics-based model used for guiding ensemble fitting is critical for accurate ensemble generation and data interpretation. Direct interpretation of specific contacts with low presence in an IDP ensemble using either all-atom or coarse-grained (CG) models still suffers from issues related to model accuracy (e.g., force field or potential energy function) and requires further experimental validation.

To understand the structural and functional relevance of transient interactions, we pursue a novel approach in this work. We focus specifically on transient interactions originating from charged amino acids, particularly between charged patches, as previous studies suggest that charge-based transient interactions can create a two-state (i.e., open and closed) behavior in a conformational ensemble.^68,69^ We simulate a series of model peptides at various temperatures and ionic strengths using a CG model to generate conformational ensembles with a broad range of transient interaction levels. Typically, the end-end distance distribution function exhibits two peaks: the first peak indicates the presence of transient interactions and suggests that the charged patches are in contact, while the second peak reflects the overall size of the protein. An adjusted polymer model enables us to quantitatively assess the fraction of transient interactions from the distance distribution function in each conformational ensemble.^69^ With this extensive database of conformational ensembles showing different levels of transient interactions, we establish an empirical relationship between sequence metrics and the fraction of transient interactions. This relationship allows us to search for transient interactions across all IDPs within the human proteome. We then explore the structural and functional relevance of these transient interactions, investigating whether they might represent a missing structural property within the sequence-structure-function paradigm for IDPs.

## Modulation of transient interactions through sequence and environmental factors

Our first goal was to establish a model system so that we could quantitatively modulate the presence of transient interactions inside an IDP. In our previous work, ^69^ we examined the fraction of transient interactions in a model peptide, DR-*N*, which comprises *N* aspartic acid residues at the C-terminal end and *N* arginine residues at the N-terminal end. The linker connecting the two charged patches was composed of glycine residues. The chain length of the peptide was set to be 100, which is adequate for analysis using polymer theories. CG simulations using the HPS model^38^ showed that the conformational ensemble of the model peptides consists of two inter-converting states: an open state where the peptide behaves similar to an IDP without any specific interactions, and a closed state where the two charged patches are in close contact with each other. This behavior is apparent in the patch-to-patch distance distribution function calculated by the distance between of the center of the two patches, as shown in Fig. 1a. The two peaks within the distance distribution function corresponds to the two primary conformational states.

**Figure 1:**
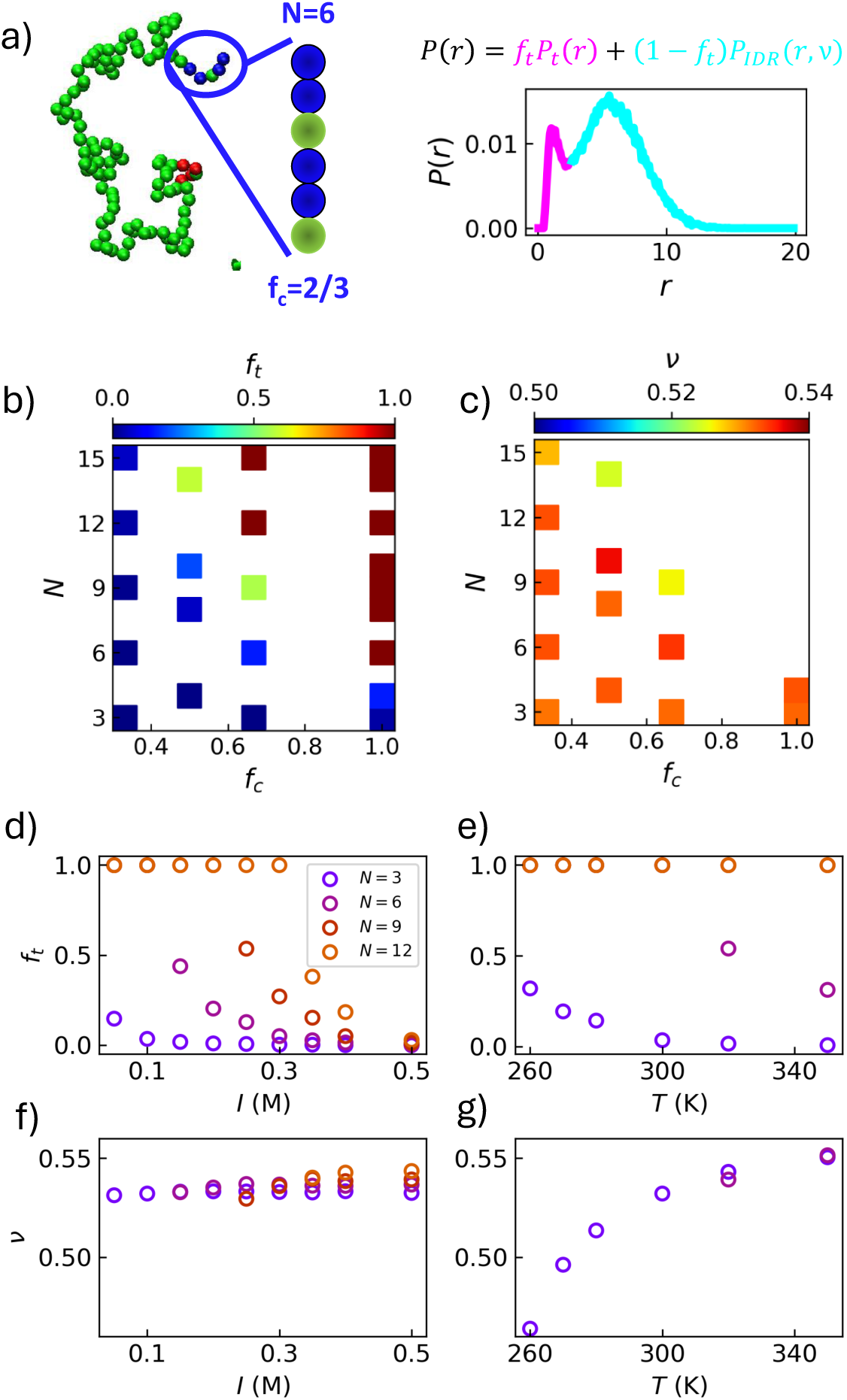
**a)** Schematic diagram of the series of model peptides characterized by two parameters, charged patch length (*N*) and fraction of charged amino acids within each patch (*f_c_*). The resulting patch-to-patch distance distribution can be modeled by an adjusted polymer model as shown in the equation, in which *f_t_* is the fraction of transient interactions and *ν* is the polymer scaling exponent. **b)** *f_t_* and **c)** *ν* values obtained from fitting Eq. 1 as a function of *N* and *f_c_* at 300 K and 0.1 M salt concentrations. **d)** *f_t_* with respect to ionic strength, *I*, at *T* =300K **e)** or temperature, *T*, at *I*=0.1 M, for multiple model peptides with different charged patch length *N* contents and *f_c_*=1 as shown in the legend. **f)** Polymer scaling exponent, *ν*, with respect to for the same cases as *f_t_* with respect to *I* **g)** and *T*. Cases where *f_t_*=1 do not display any value for *ν*.

We found that this bimodal behavior in the distribution function can be described by a combination of two distance distribution functions,^69^

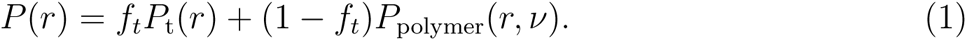

The first term is a Gaussian distribution function to match the short-distance peak,

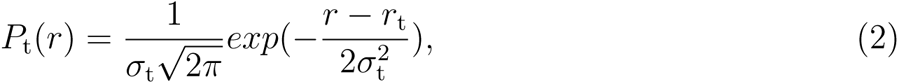

in which the parameters *r*_t_ and *σ*_t_ in the Gaussian function were fit empirically via CG simulations of a series of DR-*N* peptides to be 1.18 and 0.41 nm, respectively. The second term is a self-avoiding walk model with a varied scaling exponent *ν* (SAW-*ν* model) to match the long-distance peak. The distance distribution function of the model was initially developed to interpret the experimental measurements of an IDP in relation to its conformational properties,^52^ as shown below

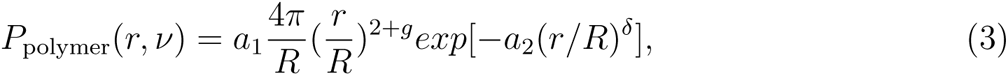

in which *g* = (*γ* − 1)*/ν* ^70^ and *γ*=1.1615 in three dimensions,^71^ *δ* = 1*/*(1 − *ν*),^72^ and *a*_1_ and *a*_2_ are normalization factors so that 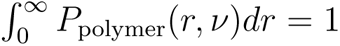 and 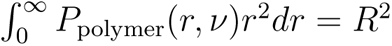. The combined model therefore contains two free parameters: the fraction of transient interactions, *f_t_*, which characterizes the weight of the short-distance peak, and the polymer scaling exponent *ν*, which characterizes the size of the peptide matching the long-distance peak. Our model reflects the metastable nature of transient interactions by considering the final *p*(*r*) function to be built from two statistically weighted parts. *f_t_* then characterizes the probability of finding the two charged patches in contact with one another and ranges from zero to one. For scenarios with low *f_t_*, the open state is still prominent in the overall ensemble. We expect that the conformational ensemble should be relatively undisturbed by the presence of transient interactions and well described by a homopolymer model, like the SAW-*ν* model.^52^ The scaling exponent *ν* for such an IDP ranges roughly from 0.45 to 0.65,^73^ serving as a valuable structural property for examining its functional correlates. However, when increasing the number of amino acids in the charge patch, *f_t_* begins to increases. The conformational ensemble of such a peptide can be thought as occupying a hybrid ensemble, switching between a homopolymer-like open state and a collapsed state. This often results in non-monotonic scaling behavior when examining the relationship between pairwise distances between amino acids and their sequence separations, indicating that *ν* alone is insufficient to fully characterize the structural properties of the IDP. For cases with large *f_t_*, the homopolymer interpretation and the *ν* value become completely irrelevant as the two charged patches will most likely be in contact.

For this work, we extended the DR-*N* model to include patches of non-consecutive charged amino acids since such a segment is more commonly seen in a biologically relevant IDP sequence. The charge content is characterized by two free parameters: the total patch length *N* and the fraction of charged amino acids within the patch *f_c_*. This allows the DR-*N* peptides to sample the sequence space in which *f_c_* is smaller than 1. To generate a series of such peptides, we generated non-consecutive patches by, for example, swapping every certain charged amino acid with a glycine residue to achieve a fraction in *f_c_*. For instance swapping every third charged amino acid will result in a new peptide with *f_c_*=2/3, as shown in Fig. 1a. In order to capture better the role of patch separation (*s*), the number of amino acids between the center of two charged patches, we also simulate peptides with different overall lengths rather than the fixed 100 residues from the previous work.^69^ This is motivated by the sequence descriptor sequence charge decoration (SCD),^20,74^ which includes sequence separation between two charged amino acids in its formula and has been shown in multiple previous works to be important to characterizing the conformational properties of IDPs.^21,36,75,76^

We first performed single-chain simulations on a series of DR-*N* peptides with different charged contents (i.e. *N* and *f_c_*) and a fixed *s*=100 over a variety of temperatures *T* and salt concentrations *I*, as shown in Table S2. The resulting data set includes 1242 combinations of sequence parameters *N* and *f_c_*, and environmental parameters *T* and *I*. In order to conduct simulations efficiently for such a large combinations of variables, we used a CG simulation model, the HPS model,^38^ which has been used to characterized the liquid-liquid phase separation (LLPS) of IDPs.^38,54^ The simulation details are included in Supporting Method A. For each case we obtained the patch-to-patch distance distribution by measuring the distance between the center of each charged patch. We then minimized the difference between the observed distribution and the adjusted polymer model via four free parameters: *f_t_* the fraction of transient interactions, *ν* the scaling exponent, *r_c_* the center of the short range Gaussian distribution, *σ_c_* representing the width of the short range peak (see Fig. S1 for fitting representative cases). We would like to note in most cases a initial guess of *f_t_*=0.5 for fitting is robust. However for extreme cases in which only one peak exists in the distance distribution function, fitting to a two-peak model is ill-defined. For such cases, we manually adjust the initial guess of *f_t_* and the bounds of minimization process to be either 0 or 1 based on the peak position. If the peak position occurs below the empirically observed cutoff distance of 2 nm, we adjust the initial guess of *f_t_* to be 1 and the allowed values of *f_t_* to be close to 1. This ensures that only short-distance function (Eq. 2) dominates the fitting. If the peak position occurs above the cutoff, we adjust the initial guess *f_t_* to be 0 and the allowed values of *f_t_* to close to 0 so that only the SAW-*ν* polymer model (Eq. 2) is present.

This process returned a comprehensive data set for *f_t_* and *ν* as a function of the sequence parameters characterizing charge patches (i.e. *N* and *f_c_*) and environmental conditions (i.e. *T* and *I*). We first looked at the fraction of transient interactions, *f_t_*, with respect to the charge content parameters. Understandably, *f_t_* approaches one for sequences with larger *N* or *f_c_* as they have higher charge content (Fig. 1b). The transition of *f_t_* from zero to one happens rapidly, occurring with the addition of only a few charged amino acids. In other words, even small variations in *N* and *f_c_* can sample the entire range of *f_t_*, indicating that *f_t_* is highly sensitive to the charge content in the sequence. This is also confirmed when checking the relation between *f_t_* and environmental parameters like *T* and *I*. *f_t_* is more responsive to variations in ionic strength than to changes in temperature (Fig. 1d and e), due to the electrostatic screening effect. However, when investigating variations in the scaling exponent *ν*, we observe a much weaker dependence, with *nu* ranging from approximately 0.5 to 0.54 across a broad spectrum of *N* and *f_c_*(Fig. 1c). This suggests that *ν* is less influenced by specific charged patches and therefore variations in ionic strength (Fig. 1f).

We would like to emphasize that for conformational ensembles where *f_t_*=1, only a single short-distance peak is present. This prevents us from determining *ν*, and scaling exponent values are not displayed for these cases. Conversely, temperature has a greater impact on *ν* values, influencing the long-distance peak and the overall size of the peptide (Fig. 1g). Since *f_t_* is primarily dependent on *N*, *f_c_* and *I* rather than *T*, it should be feasible to establish an empirical relationship between these sequence parameters and the observed *f_t_* without concern for variations in *T*.

## Empirical estimation of transient interactions driven by sequence factors

As shown previously, the fraction of transient interactions *f_t_* rapidly changes from 0 and 1 for slight variations in the sequence factors *N* and *f_c_* (Fig. 1b). We therefore propose to quantitatively model *f_t_* as a sigmoid-shaped function based on a sequence metric combining these sequence factors. From Coulomb’s law, the most obvious sequence metric would be the number of charged amino acids *q* = *f_c_N* within each of the charge patches. This way the strength of the electrostatic interaction scales with the product of *q*_+_, number of positively charged amino acids, and *q_−_*, the number of negatively charged amino acids, or here equivalently *q*^2^. We evaluated the first option by fitting the the relation between *f_t_* and *q* at different temperatures and ionic strengths using a sigmoid-shaped function, as illustrated below

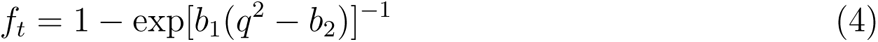

in which *b*_1_ and *b*_2_ are free parameters. A shown in Fig. S2 at three representative temperatures, we minimized the difference between Eq. 4 and the values of *f_t_* obtained from simulations described in the previous section. The fitting works reasonably well for extreme values of *f_t_*=0 or 1 but we see clear deviations in the transition region when *f_t_* varies rapidly.

Quantifying the strength of electrostatic interactions using *q* assumes that all oppositely charged amino acids are always in contact without obstruction through the simulations. This might be possible for a fully rigid rod, however this is not the case for an IDP since the charged patches exhibit flexible conformations. The effective number of charged amino acids contributing to the attractive electrostatic interactions might be smaller than the real number of charged amino acids *q* within the patch. To improve the fitting shown in Fig. S2, we need to introduce a correction on *q*. Since the deviation comes from the flexible conformation of the charge patch, we borrowed an idea from the scaling concept in the polymer theory by introducing a scaling exponent *β* on *N* when calculating the effective number of charged amino acids, as shown in the equation below

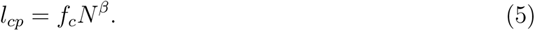

This new metric will be referred to as the effective charge patch length (*l_cp_*). We performed a global fitting of the *f_t_* curves for all simulation results, replacing *q* with 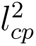, as

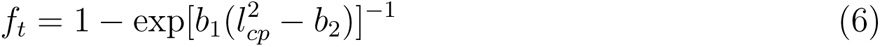

In terms of the free parameters in the fitting, *b*_1_ and *b*_2_ in Eq. 5 are permitted to vary independently for each curve, but the *β* value are kept the same across all curves. This global fitting returned an optimal scaling parameter of *β* = 0.76. The representative fitting curves for simulations at 300K are shown in Fig. 2a and the curves for simulations at all other temperatures are shown in Fig. S3. Even for large salinity (i.e. 0.5*M*), in which transient interactions did not form, the sigmoid function can still capture the variation of *f_t_*. This demonstrates that a single *β* value is sufficient to fit all *f_t_* curves for any environmental conditions. An optimal *β* value of 0.76, compared to 1 in the case of *q* (Fig. S2), suggests the conformation of the charge patch is less rigid than a rod but more rigid than a random coil with a scaling exponent of 0.5. This may by due to electrostatic repulsion within the patch. One remaining question before using the new sequence charge metric *l_cp_* is whether *β* is robust enough to varying environmental factors. We found that *β* is minimally affected by ionic strength if we fit the simulations of multiple sequences with independent *β* values at different ionic strengths and the same temperature. This is as expected since ionic strength only affects the electrostatic interactions that can be quantified by *l_cp_*. However when applying the same test by fitting the data independently at each temperature as shown in Fig. 2b, we did observe slight variation of *β* at different temperatures. This is mainly due to the impact of temperature on the overall size or *ν* of the peptide, which will slightly affect the balance between the short- and long-distance peaks in the patch-to-patch distance distributions (Fig. 1a). Since varying *β* does not significantly improve the fitting, and the optimal *β* for each temperature falls within a narrow range of approximately 0.7 to 0.8, we have chosen to use a constant *β*=0.76 for all temperatures moving forward.

**Figure 2:**
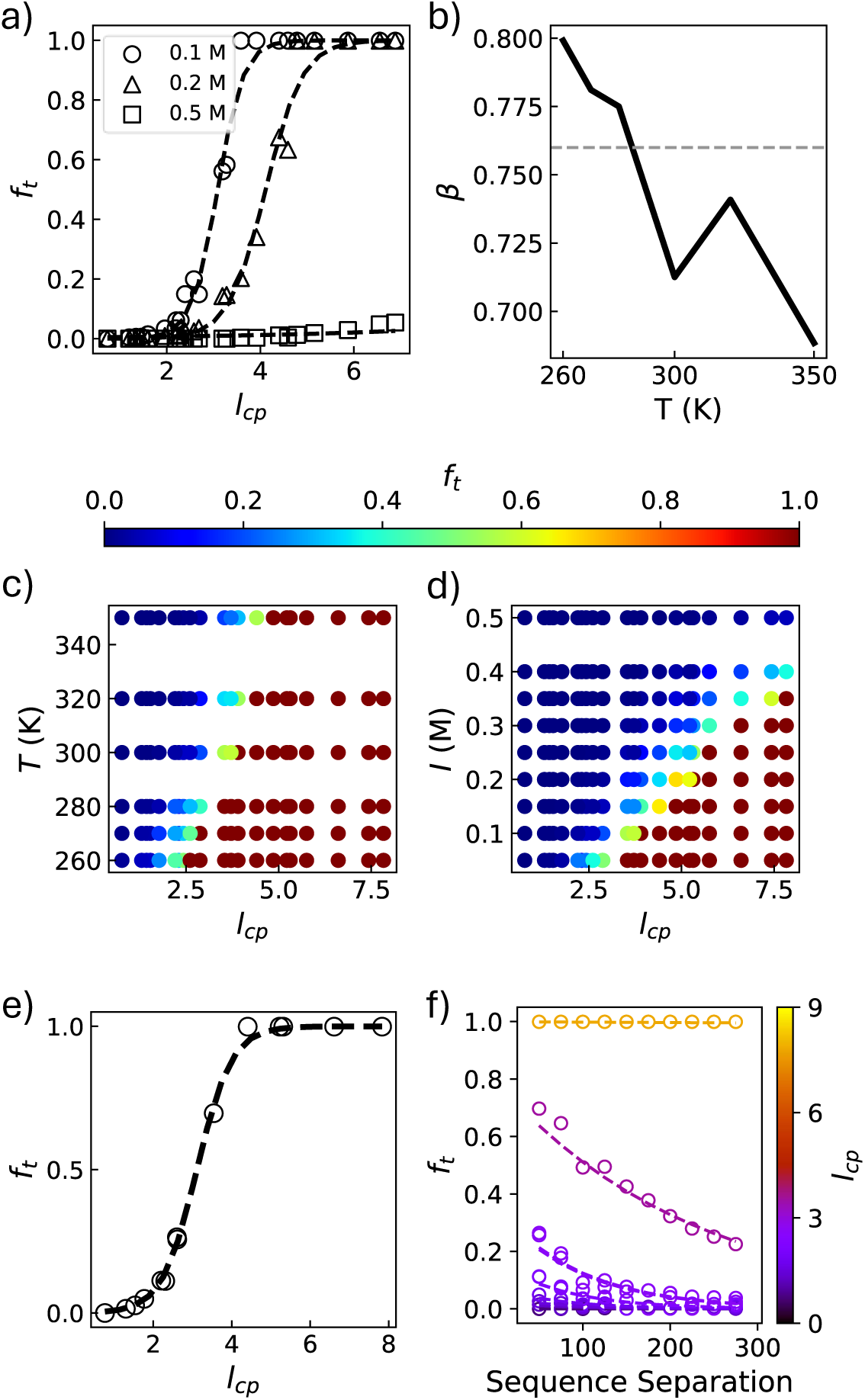
**a)** Effective charge patch length (*l_cp_* = *f_c_N^β^*) underlies *f_t_* in a sigmoid-shaped function. Open shapes indicate data from simulations while dashed lines represent the global fitting to Eq. 6. The ionic strength *I* is shown in the figure legend. **b)** *β* as a function of *T* obtained by fitting data at individual *T* independently. The dashed line indicate the *β* value determined from a global fitting of data at all *T*. **c)** *f_t_* as a function of *l_cp_* and *T*. **d)** *f_t_* as a function of *l_cp_* and *I*. **e)** *f_t_* as a function of *l_cp_* at a patch separation of *s*=30. Open circles indicate simulated data, while the dashed line displays best fit results using Eq. 6. **f)** *f_t_* as a function of *s* for sequences with different *l_cp_*. Open dots come from simulations, while dashed lines indicate fitting results from Eq. 7.

We further investigated the role of charge content on transient interactions using the new sequence charge metric *l_cp_*. This metric effectively consolidates the two original sequence metrics (*N* and *f_c_*) into a single parameter. Thus we can simultaneously monitor three different variables: *l_cp_*, *T* and *f_t_* (Fig. 2c), or *l_cp_*, *I* and *f_t_* (Fig. 2d). In both figures that include *T* and *I*, we observed a rapid transition of *f_t_* as a function *l_cp_*, which aligns with our assumption of employing a sigmoid-shaped function (Eq. 4) to capture this behavior. Across the range of temperatures we simulated, the point at which *f_t_* begins to transition shifts from roughly *l_cp_*=2 to *l_cp_*=4. In contrast, *I* induces a very significant shift in the onset of transient interactions, changing from roughly 2 to 7.5. A more quantitative analysis of the threshold *l_cp_* values that drive certain *f_t_* at different *T* and *I* is presented in Fig. S4. Ionic strength emerges as a more significant environmental factor, influencing both the onset and speed at which *f_t_* transitions. *T* meanwhile primarily regulates the overall size of the peptide and has a limited influence on *f_t_*. Notably, we discovered that a small threshold *l_cp_* value of approximately 3 is sufficient to achieve 50% of transient interactions at 300K and 0.1M, corresponding to a modest charged patch consisting of three consecutive charged amino acids. Furthermore, the required threshold *l_cp_* values are sensitive to small variations in *I*, suggesting that *f_t_* can be fine-tuned through minor sequence adjustments.

The other sequence metric that is expected to influence the prevalence of transient interactions is the patch separation *s*, defined as the number of amino acids between the centers of the two charged patches. To characterize this, we conducted additional simulations at a fixed temperature of 300K and an ionic strength of 0.1M, varying *s* from 30 to 300 residues (Table S3). The *f_t_* values were analyzed by the same method as shown in the previous section. It is important to note that obtaining *f_t_* from the simulated *P* (*r*) at extremely low patch separations is not feasible, as the peak representing the open state significantly overlaps with the peak corresponding to the closed state in the patch-patch distance distribution function. This overlap complicates the fitting process, making it challenging to determine *f_t_* and *ν* simultaneously using Eq. 1. Therefore the smallest *s* in our database is 30. As shown in Fig. 2e for *s*=30 and in Fig. S5 for all the other *s*, the globally determined scaling value *β*=0.76 can still be utilized to fit *f_t_* for these new sequences. We observed that, for a fixed value of *l_cp_*, *f_t_* decreases as a function of *s*. This trend holds true only for sequences where *f_t_* is less than one; otherwise, transient interactions remain robust across the range of *s* that we investigated. This indicates that beyond a certain level of charge content, it becomes virtually impossible for contacts between the two patches to break once formed, resembling the stable native contacts found in folded proteins. Motivated by this observation, we introduce an exponential function to fit the relation between *f_t_* and *s*, as shown below

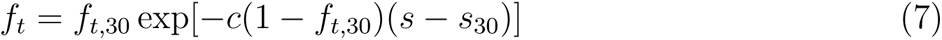

in which *c* is a shared free parameter for simulations at all *s*, *s*_30_=30 is our reference patch separation and *f_t,_*_30_ is the fraction of transient interactions at *s*=30 as shown in Fig. 2e. Eq. 7 recapitulates *f_t_* for simulations at a wide range of patch separations tested, seen in Fig 2f. Together with Eq. 6, whose free parameters were obtained by fitting *f_t_* for *s*=30, *f_t_* for any sequences can be predicted by these two empirical equations using the two sequence metrics *l_cp_* and *s*.

At this point, we have identified several key principles regarding the fraction of transient interactions *f_t_* driven by charged amino acids. First, *f_t_* can be characterized by an effective charge patch length *l_cp_* with a scaling parameter less than one to the charge patch length. Second, transient interactions are diminished by increasing ionic strength and electrostatic screening. Third, transient interactions become slightly less favorable with raising temperature due to the overall increase in peptide size. Fourth, *f_t_* decay exponentially with patch separation *s*. And finally, *f_t_* can be quantitatively predicted using the empirically derived Eqs. 4 and 7.

## Influence of transient interactions on conformational behaviors

With the sequence sources of *f_t_* thus characterized, we moved on to understand the effect these transient interactions have on single- and multi-chain structural behaviors. We introduced two additional classes of simulations: slab coexistence and cubic box simulations. The slab simulation protocol is described in Supporting Method B and in a previous work.^38^ All simulations were performed at 280 K, slightly below the critical temperature for most DR-*N*. This temperature was selected in order to observe co-existence of the two phases, across the same range of ionic strengths as the single-chain simulations described in the previous section. The sequence and environmental factors of all the slab simulations are shown in Table S4. In order to further understand the conformational behaviors of the peptides in the high-density phase, we performed additional cubic box simulations at a concentration slightly larger than concentration of the high-density phase (Table S5). Such cubic box simulations reduce the complexity of assigning chains within and without of the slab during the analysis. These new simulations allowed us to characterize the effect of transient interactions on phase separation, and more interestingly, the inner structural behaviors of the resulting liquid droplets. For illustrations of structural observables in this section, we use a unified color code along the new sequence metric *l_cp_* introduced in the previous section, which correlates with the fraction of transient interactions *f_t_*, as shown in Fig. 3a.

**Figure 3:**
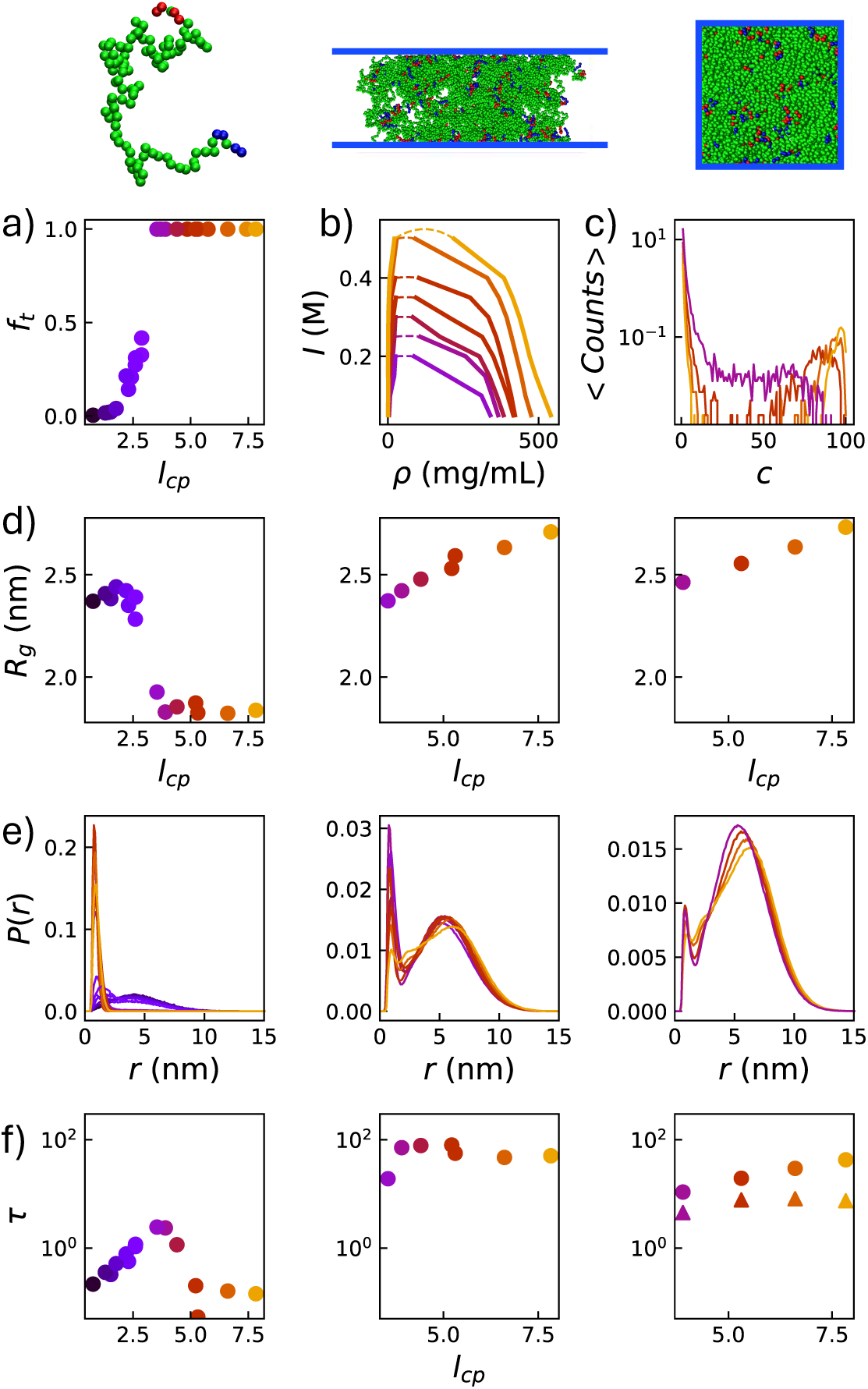
Snapshots from each of the three simulations types (single-chain, coexistence slab, cubic box), with the corresponding information for each type of simulation aligned in the respective columns. Blue lines indicate the positions of the periodic box. **a)** *f_t_* as a function of *l_cp_* for *T* = 280 K and *I* = 0.1 M. *l_cp_* is represented by the same color code in the remaining subfigures. **b)** *I*-dependent phase diagrams from slab coexistence simulations. **c)** Histogram of number of peptides in each cluster per frame from cubic box simulations. **d)** Radius of gyration for each simulation case. **e)** Intra-chain patch-patch distance distribution function in each simulation type. **f)** Relaxation time for the three simulation classes. Circular marks display the relaxation time from intra-chain patch-patch distances while triangular marks indicate the relaxation time from inter-chain coordination numbers.

First, we obtained the density profiles from the coexistence slab simulations as shown in Fig. S6. Only sequences with *l_cp_* greater than values ∼ 3.8 formed stable condensates at *T* =280K and *I*=0.1M, in which two phases coexist. The concentrations of the two phases will be further referred to as the saturation concentration, which is the concentration of the low-density phase and the threshold concentration to observe LLPS, and the dense phase concentration, which is the protein concentration inside the liquid droplet. We plotted these two concentrations at different *I* onto a phase diagram, as shown in Fig. 3b. Interestingly, *f_t_* is already fully saturated for these sequences in single-chain simulations, seen in Fig. 3a, while dense phase concentration continues to increase with *l_cp_*. The critical ionic strength *I_c_*, below which LLPS is possible, can then be obtained by fitting the phase diagram via the equations shown in Supporting Method B. As shown in Fig. S7, *I_c_* and the dense phase concentration increase similarly when increasing *l_cp_* whereas the saturation concentration drops. Such behavior appears to be a result of more favorable interactions towards phase separation. While higher *l_cp_* and charge content contribute to phase separation, electrostatic screening due to higher ionic strength effectively reduces interactions between charged amino acids and therefore, above a certain threshold, converts the two phases into one.

Our next task was to see whether any microscopic structures exist within the condensate, such as a network connecting multiple peptides. This phenomenon is often referred to as percolation coupled phase separation.^65,77^ The analysis of the formation of the peptide clusters cannot be easily applied to a coexistence slab simulation due to existence of two phases. We therefore followed a framework previously introduced by Das et. al.^66^ by analyzing peptides clusters in a cubic box simulation. A cluster consists of all peptides that share common contacts within their oppositely charged patches (see Supporting Methods C). For example, if two chains have at least one pair of residues from their oppositely charged patches within a certain cutoff distance and a third chain is also in contact with one of them, then all the three belong to the same cluster. For each frame of a simulation, we identify multiple clusters containing a certain number of peptides *C*, and obtain the distribution of *C* throughout the simulation, shown in (Fig. 3c) using a cutoff distance of 0.7 nm. This process was tested using a range of contact cutoff distances, as displayed in Fig. S8, and qualitatively similar results were observed. For low *l_cp_* values, we observed a broad distribution of cluster sizes although the largest peak occurs close to *C* = 1, which represents chains without a cluster. This indicates that connections are forming but specific transient interactions are not strong enough to bridge the entire condensate. For higher *l_cp_* sequences, the population of the middle *C*-value region disappears. Here, charge interactions become strong enough to connect most chains in the system, although unclustered chains still remain. This might be due to the similarly increasing intra-chain interactions which make it difficult for a chain to join the condensate network. This behavior is consistent with previous publications ^65,66,77^ that show how multivalent interactions between peptides lead to such microscopic network structure and therefore percolation coupled phase separation within the condensate. Here in our model, these multivalent interactions come from specific transient interactions between charged patches and are well characterized by *l_cp_*.

We would like to further explore how other structural features correlate with *l_cp_* and consequently the network structures within the condensates. Intuitively, these structural features should be primarily controlled by charge-patch-driven intra- and/or inter-chain interactions. To see if this was the case, we investigated the properties of individual peptides from monomer simulations to those in the dense phase beginning with radius of gyration (*R_g_*). In the single chain case, *R_g_* is largely shaped by the presence of transient interactions. As shown in Fig 3d, *R_g_* collapses with respect to *l_cp_* as *f_t_* saturates. This observation is straightforward, as peptides sampling only the closed conformation will have a lower *R_g_*. In the slab and box cases, we realized two interesting observations. The first observation is that *R_g_* is significantly higher and become less sensitive to the variations in *l_cp_* compared to the single chain cases. This behavior, where *R_g_* is less affected by environmental conditions or amino acid sequences in the condensate, has been previously reported both computationally^78,79^ and experimentally,^80^ as peptides strive to maximize favorable interactions with one another.

However, the second observation that *R_g_* even slightly increases and scales linearly with *l_cp_* is counter intuitive. We interrogated this possibility by first obtaining patch-to-patch distance distributions from each case. Single chain distributions (Fig 3e) *P* (*r*) shift from the open, polymer-like state to the closed state with increasing charge content. Corresponding distance distributions within the condensate in slab and cubic box simulations do just the opposite. In less dense condensates (*l_cp_*∼4), the closed state is still prevalent, although these may be attributed to chains in the dilute phase. For the highest values of *l_cp_*, the closed state disappears almost entirely and gives way to a large open state peak. However, this open state differs from that observed in the single-chain case, as the width of this peak ranges from approximately 4 to 12 nm. This suggests that, while patch-to-patch contacts are forming between peptides, they do not necessarily constrain the ends of each peptide within a narrow range. We further explored the the relation between inter- and intrachain interactions raised by charged amino acids, movtivated by two recent work.^78,81^ Intra- and inter-chain contacts are calculated as the average number of positive to negative patch contacts per frame, where contacts are determined using the same criteria as in the clustering analysis (Fig. S8). Regardless of the cutoff distance applied, contact propensities behave qualitatively the same with respect to *l_cp_*. Not just intra- but also inter-chain contacts increase linearly with respect to charge content, roughly doubling over the range of *l_cp_* values considered. Thus, in addition to promoting the formation of large, condensate-spanning clusters observed previously, stronger inter-chain contacts contribute to a more expanded *R_g_* that is less influenced by environmental conditions or amino acid sequence variations.

Finally we performed two types of dynamics analysis (see Supporting Method D) to quantify the rigidity of the condensate network structure. Intra-chain distance correlation was calculated for chains within the slab and compared to those for single chains, shown in Fig. 3f. For freely moving chains, the relaxation time *τ* is controlled by the transition between the open and closed state. This is evident from the large peak in *τ* as a function of *l_cp_* for the region where *f_t_* is close to 50%. For sequences where one state dominates (i.e. *f_t_*=0 or 1), *τ* is contributed by the relaxation within each conformational state and is therefore smaller than the *τ* for the transition between the two states. Intra-chain motions relax much slower for chains within a condensate in both co-existence slab simulation and in the cubic box simulation. Even for the least dense slab, *τ* is two fold larger than the maximum single-chain value. Increasing *l_cp_* slows down intra-chain relaxation due to the increasing population of the open-state conformations and slow relaxation of the transitions between the two states. Evidently, intra-chain motions are not fully restrained even for the largest *l_cp_* values visited in our simulation. We further investigated the impact of *l_cp_* on the inter-chain relaxation. Since one charge patch might interact with several other oppositely charged patches at the same time, it is not possible to simply analyze the relaxation of the inter-chain distance between one specific pair of charge patches. We therefore calculated the coordination number for each charge patch, namely the number of charge patches in contact with the charge patch of interest, as a function of the simulation time. The two charged patches are considered in contact when the distance of at least one pair of residues is smaller than 0.7 nm, just as with the clustering and contact analysis cases. The coordination number here is clearly a feature of the formation of nodes, which comprise the network structure within the condensates. A similar relaxation measure can then be applied to this coordination number, as displayed in Fig. 3f. *τ* in this case increases linearly with *l_cp_* similar to the intra-chain relaxation. However *τ* from inter-chain relaxation is one order of magnitude faster than that from intra-chain relaxation. This suggests two notable structural behaviors of the network structure: 1) robust nodes that exhibit fast relaxation in terms of the number of peptides forming the node; and 2) flexible linkers with slow relaxation, as the two ends are partially restrained by the nodes.

## Exploration of the functional relevance of transient interactions

As we have demonstrated, the presence of transient interactions has a significant impact on the conformational behaviors of a model peptide. Our next question was to see whether any correlation exists between the fraction of transient interactions within an IDP and its function. In other words, do certain classes of IDPs rely on specific transient interactions to perform their roles? We addressed this question in two steps: first by adapting our previous method to estimate the fraction of transient interactions *f_t_* to IDP sequences from a biological database and second by performing Gene Ontology (GO) enrichment analysis^82^ for biological sequences with similar *f_t_* values.

In the previous section, we showed that *f_t_* driven by two charge patches can be estimated using Eqs. 4 and 7, which contain two sequence dependent input parameters: *l_cp_* the effective charge patch length and *s* the patch separation. However, biological sequences might not have well defined “charge patch” areas for us to directly calculate *l_cp_* and *s*. To adapt our previous method to estimate *f_t_* from any arbitrary sequence, we introduced a patch scanning algorithm to determine the optimal patch length *N* and dominant positive and negative charge patches within the sequence. First, we scanned a range of window sizes from 3 to 15, the same range tested in our model peptide. *l_cp_* was calculated for each window using Eq. 5, where *N* is then the window size and *f_c_* is the net charge per residue instead of fraction of charged amino acids. This allowed us to account for the coexistence of both types of charged amino acids within the patch. For each sequence, we obtained *l_cp_* as a function of both sequence position and patch size, shown in Fig. 4a using the disordered part of S13A4 as an example. Second, we retained the smallest, most negative *l_cp_* value as *l_cp−_* and the largest, most positive as *l_cp_*_+_, as well as their sequence separation *s*. These values correspond to the largest effective negative and positive charge patch length (shaded area in Fig. 4a), respectively. In order to avoid selecting overlapping positive and negative patches which might exist in sequence with well-mixed charges, we checked if the identified patches overlap with one another and removed overlapping patches from the *f_t_* determination. At last we estimated *f_t_* using Eqs. 4 and 7 substituting 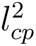 with the product of *l_cp_*_+_ and *l_cp−_*. The estimated *f_t_* for the protein S13A4 is 0.36 for the two patches with a patch separation of 86 amino acids. The smallest sequence separation *s* between the two charged patches we visited is 30 since we are more interested in nonlocal transient interactions.

**Figure 4:**
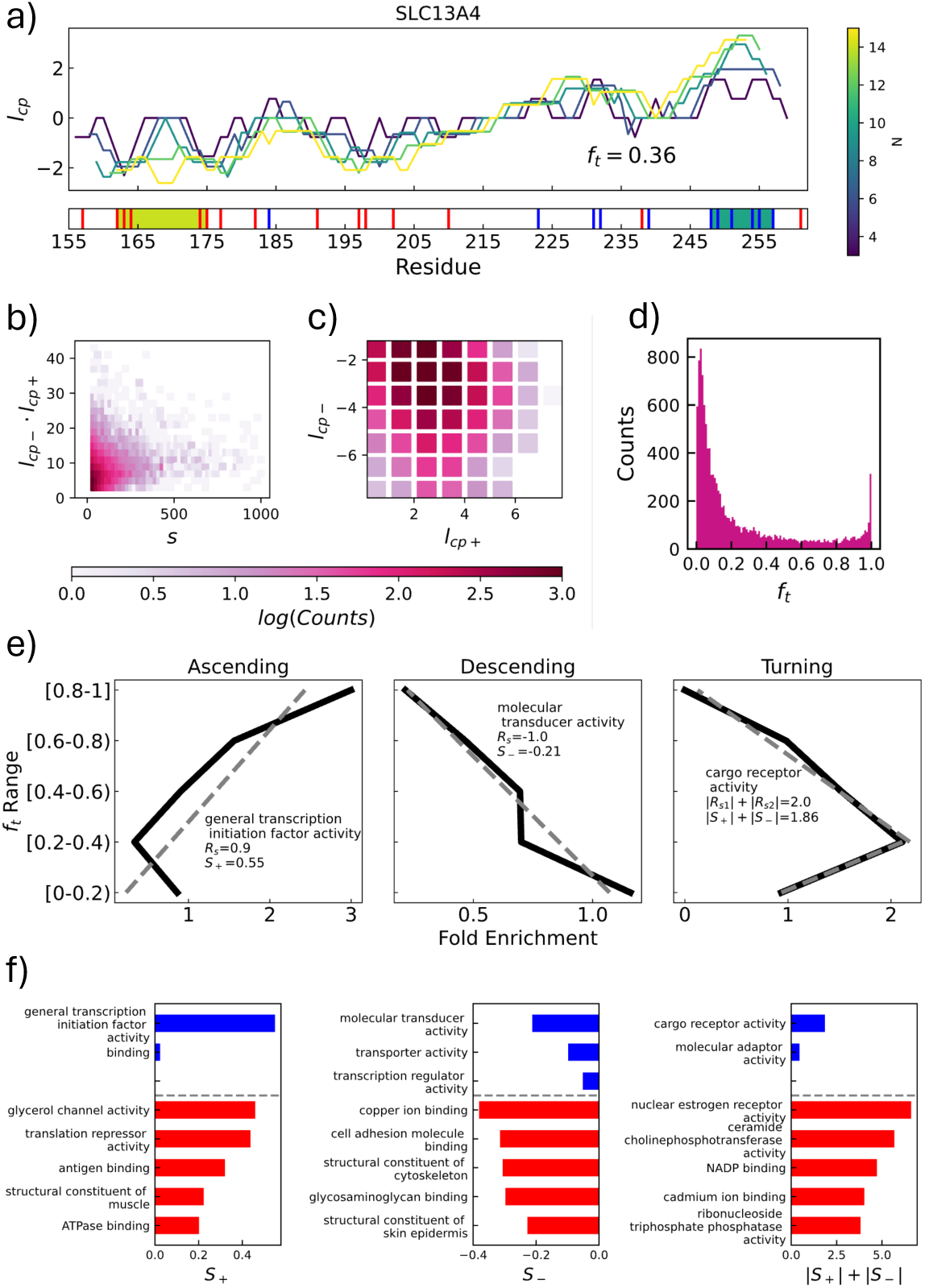
**a)** Schematic diagram of the *l_cp_* calculating algorithm using the disordered region of the SLC13A4 protein as an example. Positive charges are denoted along the sequence in blue while negative charges are denoted in red. Shaded regions indicate the size and location of the strongest positively and negatively charged patches determined from the algorithm. **b)** Histograms of the product of positive and negative effective charge patch length (*l_cp_*_+_ *l_cp−_*) and patch sequence separation *s*, determined by the scanning algorithm throughout the IDRome database.^23^ **c)** Histogram of *l_cp_*_+_ and *l_cp−_*. **d)** Histogram of *f_t_* values predicted. **e)** Three representative GO terms categorized via the trend between fold enrichment and *f_t_*. The dashed lines indicate the results from a linear fit. Legends show the GO terms, Spearman correlation coefficients (*R_s_*) and positive and negative slopes (*S*_+_ and *S_−_*) from linear fit. For the turning case, Spearman correlation coefficients and and linear fitting are applied to both sides of the fold enrichment peak. **f)** Slope from linear regression for high (blue) and low-level (red) GO terms for each category.

Admittedly, this scanning algorithm has several caveats. First, sequences may contain some charged residues outside the identified patches, and smaller charge patches may also contribute to determining *f_t_*. Due to the strong sigmoid dependence between *f_t_* and *l_cp_*, these smaller patches might play a secondary role on *f_t_* (Fig. 2a). Second, evidence suggests that when the same type of charged amino acids segregate into a short fragment, their effective charge might be smaller than the total number of charged amino acids due to charge regulation and/or renormalization.^83,84^ In such cases, our algorithm tends to overestimate *f_t_*. However, this is unlikely to affect the next step of checking the qualitative and not quantitative functional relevance of transient interactions.

To apply our *f_t_* determination algorithm, we turned to the IDRome database,^23^ which contains more than 28000 disordered sequences. We first performed our *l_cp_* scanning algorithm and identified approximately 11000 sequences with a pair of non overlapping positively and negatively charged patches. As shown in Fig. 4b, patch separations *s* can be as large as approximately 1000 residues, although the majority of patch separations are shorter than 400. Values of *l_cp,_*_+_ and *l_cp,−_* are not equally distributed and strong negative patches are slightly more common than positive ones (Fig. 4c). Feeding the product of *l_cp,_*_+_ and *l_cp,−_* and patch separation *s* into Eqs. 4 and 7, we estimated *f_t_* for the identified patch pairs. We then binned the sequences according to the *f_t_* values, as shown in Fig 4d. Most IDPs (approximately 84%) exhibit little to no transient interactions (0 *< f_t_ <* 0.2), consistent with the traditional understanding that IDPs lack specific interactions. However, approximately 11% of IDPs show *f_t_* values somewhere between 0.2 and 0.8, suggesting that transient interactions are not rare in IDPs. Interesting, we see plenty of sequences (approximately % 5) with an *f_t_* value above 0.8, suggesting almost stable specific interactions. These interactions, often seen in sequences with high fraction of charged amino acids, cause the conformational ensemble to predominantly sample the closed state. They might only be seen as transient interactions for high ionic strength and strong electrostatic screening.

To determine which biological functions have a dependence on transient interactions, we performed GO enrichment analysis on the collection of sequences based on their *f_t_* values. First we classified IDP sequences into five groups based on their *f_t_* value ranges and collected their GO terms. Due to the tree structure of GO terms, we scanned the list of molecular function GO terms for each sequence and always added back the corresponding parent terms. Each sequence therefore contains on average four to eight GO terms, varying betweenm different *f_t_* ranges (see Fig. S9). With this finalized, we calculated each term’s fold enrichment throughout the *f_t_* groups with respect to the whole IDRome database as *N*_grp_*T*_ref_ */*(*N*_ref_ *T*_grp_), in which *N*_grp_ is the number of times a term is annotated within a group, *N*_ref_ is the number of times a term appear within the reference database, and *T*_grp_ and *T*_ref_ are the sizes of the *f_t_* group and reference set, respectively. Each term therefore has a series of five values representing its fold enrichment in different *f_t_* ranges as shown in Fig. 4e.

We used these values to create a metric which separated the GO terms into four classes based on their dependence with respect to *f_t_*. Since sampling GO terms within a specific *f_t_* range might be limited, we only classified terms that are represented in at least four out of the five *f_t_* ranges. First we calculated the Spearman correlation coefficient to determine if the term’s fold enrichment monotonically increases or decreases with respect to *f_t_*. Terms whose correlation coefficient is great than 0.8 or smaller than −0.8 fall into one of the first two classes, “ascending” or “descending” (Fig. 4e). The strength of the dependence between the GO term and *f_t_* can be quantified by the linear fitting slope *S*_+_ and *S_−_* for these two classes, respectively. Terms which do not meet this criteria are subject to a second test. We obtained the maximum value of fold enrichment within different *f_t_* ranges and again calculated the Spearman correlation coefficient on the two sides of the maximum. This test determines which terms have a non-monotonic behavior with respect to *f_t_*. Terms whose first and second Spearman coefficients are above 0.8 and below −0.8 fall into the final class, “turning”. The strength of the dependence between the GO terms and *f_t_* in this class can be quantified by |*S*_+_| + |*S_−_*|, where *S*_+_ and *S_−_* represent the linear fitting slopes from either side of the maximum. All others GO terms which were not classified into the previous classes were sorted into the “none” class. These classifications provide us a guideline for determining the preference of specific GO terms to certain levels of *f_t_*. Ascending means that the specific GO term tends to sample sequence with high *f_t_*, descending low *f_t_* and turning medium *f_t_*. The “none” class suggests unclear dependence of fold enrichment values across different *f_t_* ranges.

We concentrated on two distinct categories of GO terms in our analysis: 1) the highest-level terms, which are the immediate child terms of the broader category molecular function, and 2) the lowest-level child terms, which represent the terminal vertices in the GO tree map. These will henceforth be referred to as high-level and low-level terms, respectively. The high-level terms correspond to general molecular functions that are parent nodes in the hierarchy, while the low-level terms represent the most specific, finely categorized functions that do not branch further. This distinction allows us to analyze biological processes from both a broad perspective and a detailed, function-specific one, providing insights into how the conformational preferences of IDPs might correlate with different functional roles at varying levels of specificity. As shown in Fig. S9, we observed that the high-level terms exhibit a roughly equal distribution across the three classes—ascending, descending, and turning, representing the preferences of transient interactions. This suggests that, at a broader functional level, all three scenarios of transient interactions might be useful in tuning biological processes. On the other hand, the low-level terms showed a marked preference for the turning class, which refers to a particular dynamic conformational ensemble of IDPs in which the protein undergoes rapid structural adjustments due to medium transient interactions. This suggests, at more specific functional levels, certain IDP sequences may rely heavily on this flexible, transient state to carry out precise molecular functions. Additionally, we sorted the GO terms in three classes according to their linear fitting slopes *S*_+_, *S_−_* and |*S*_+_| + |*S_−_*|, respectively, and showed the representative high- and low-level terms in Fig. 4f. While these data do not serve as direct evidence that specific functions require the presence of transient interactions, we do observe several interesting correlates. For instance, in the ascending class, the general transcription initiation factor stands out as the highest-ranking term. Proteins within this category are known to mediate the assembly of the RNA polymerase holoenzyme and contain clusters of charged amino acids.^85^ In the turning class, nuclear estrogen receptor activity is the highest-ranking term. Notably, one member of this class has recently been shown to exhibit functionally relevant transient interactions.^28^ These findings highlight the potential role of medium-strength transient interactions in regulating molecular processes across different contexts.

## Conclusion

In this work, we have investigated how patches of charged amino acids contribute to predictable levels of specific, transient interactions. Based on a series of model peptides with different charged patches, we established a large database of transient interaction levels under varying sequence and environmental conditions using coarse-grained simulations. The level of these transient interactions can be reliably modeled by a polymer model and predicted by a new sequence metric, the effective charge patch length.

Using this new sequence ruler for transient interactions, we further explored the impact of different levels of transient interactions on the structural and functional behaviors of IDPs. Notably, we observed increasing network structures during liquid-liquid phase separation as transient interactions intensified. This phenomenon is primarily attributed to the conversion from intramolecular to intermolecular transient interactions between charged patches within the condensates. Additionally, we noted a significant enrichment of molecular functions associated with medium-strength transient interactions. This highlights the potential functional relevance of these dynamic interactions, which are neither too weak, converting an IDP into a random coil, nor too strong, causing it to adopt a folded conformation. This delicate dynamic balance of transient interactions may be crucial for enabling IDPs to adopt multiple conformations and fulfill diverse biological roles.

While transient, specific interactions are undoubtedly significant structural features among some IDPs, our description of these interactions is clearly incomplete. First, transient interactions can arise from sequence characteristics beyond just charged amino acids; they can also be influenced by hydrophobic and aromatic residues. Additionally, IDPs often engage with various interaction partners, suggesting that specific interactions may play a role in heterotypic interactions as well. Our analysis of transient interaction enrichment in the human proteome also lacks a mechanistic understanding of why charge-based interactions are essential for realizing specific functions. Nevertheless, our initial evidence indicating that more than 10% of IDPs exhibit charge-driven transient interactions strongly suggests that this phenomenon is not uncommon and deserves further exploration. Continued research in this area will hopefully illuminate more missing structural elements within the IDP sequence-structure-function paradigm.

## Acknowledgement

This work is supported by the National Institutes of Health (R35GM146814). The authors acknowledge Research Computing at Arizona State University.

## Supporting Information

### 1. Supporting Methods

#### Coarse-grained (CG) model

The CG HPS model[1] considers each amino acid as one bead, described by three types of interactions: bonded, electrostatic and short-range pairwise interaction terms characterized by amino acid hydropathy.[2] Bonded interactions are modeled by a harmonic potential with a spring constant of 10 kJ/Å^2^ and a bond length of 3.8 Å. Electrostatic interactions are modeled using a Coulombic term with Debye-Hückel [3] electrostatic screening,

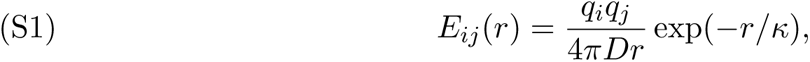

in which *κ* is the Debye screening length and *D* = 80, the dielectric constant of the solvent. The short-range pairwise interaction is modeled using Ashbaugh-Hatch functional form[4],

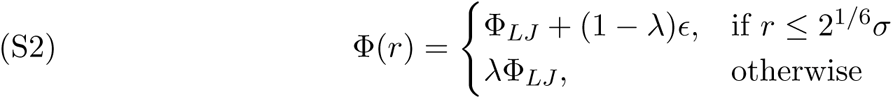

in which Φ*_LJ_* is the standard Lennard-Jones potential

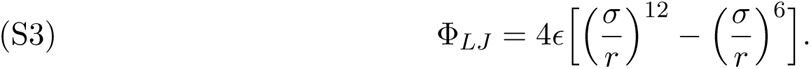

The *λ* value in the pairwise interaction term is the arithmetic average of the *λ* values of the two corresponding amino acids. The amino-acid specific parameters of the model are shown in Table S1. The interaction strength *E* is set to 0.2 kcal/mol based on parameterization from previous works.[1]

The HOOMD-Blue software v2.9.3 [5] together with the azplugins [6] were used for running the molecular dynamics simulations. All simulations were run using a Langevin thermostat with a friction coefficient of 0.01 ps*^−^*^1^, a time step of 10 fs and a temperature of 298 K for 5 *µ*s. The simulations were recorded at every 100 ns, which resulted in 50000 conformations for each condition. The first 1000 frames (0.5 *µ*s) were dumped for equilibration and the remaining 49000 frames were used for further analysis. Distance distributions are calculated for every simulated CG trajectory by recording the distance between the center of the two charged patches. The histogram was made using 100 bins and a distance range of 0 to 20 nm.

#### Slab simulation method

Slab coexistence simulations were conducted in three steps. First, a large cubic box was populated with 100 peptide chains. The simulation box was then gradually compressed over 10 ns to achieve a high peptide concentration while avoiding steric overlaps. In the final step, the simulation box was extended along the z-axis to 280 nm, significantly larger than the x- and y-axes. This configuration created a slab-like structure in the center of the simulation box, effectively mimicking a droplet with an infinite surface area. All slab simulations were then run for 5 *µ*s, with the first 1 *µ*s designated for equilibration, and data analysis performed on the remaining 4 *µ*s trajectories.

To determine the dense phase and saturation concentration of a given slab simulation, we first centered the slab of each frame to *z*=0. We then calculated the density profiles with 100 bins averaged over the last 4 *µ*s (40000 frames) of the simulation. The dense phase concentration is then given by the average density at *z*=0, the center of each slab, while the saturation concentration is calculated by the average of the 10 bins furthest from the center. After obtaining both concentrations for all sequences and simulation conditions, we discarded any cases where the difference between saturation and dense phase concentration was below a certain cutoff. The cutoff (approximately 25 mg/mL) was established by estimating the low-density and high-density phase concentrations from the slab with the lowest charge content, *N* = 3*, f_c_* = 1*/*3, which was determined by inspection not to form a condensate.

Critical ionic strength was determined by fitting the remaining data to Eq. S4 with two free parameters, *A* and *I_c_*, the critical ionic strength,

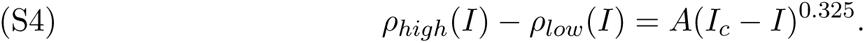

In order to achieve a meaningful fitting, we only considered sequences for which phase separation was observed in at least three different ionic strengths. In other words, any slabs which disassociated too quickly with respect to ionic strength were discarded.

#### Cubic box simulations

Cubic box simulations were performed so that each box size created an environment at a slightly higher concentration than that of the high-density phase. This ensured that the box consisted of only one phase to calculate relevant structural features. This was achieved by calculating the high density phase concentration for each sequence at 0.1 M salt concentration and then solving for the size of cubic box which would contain 100 chains at the same concentration. The actual simulated box was 0.1 nm short on all sides than the solved case.

For each frame of the cubic box simulations, we applied a cluster-searching algorithm to identify peptide clusters. Starting with an initial chain, we identified all other chains in contact with it based on interactions between their positive and negative patches, where contact was defined as at least one residue from a positive patch contacting a residue from a negative patch on the chain of interest. Various distance cutoffs for these contacts were tested (Fig. S8). For each contacted chain, the same search process was iteratively applied to find additional contacting chains, continuing until no new contacts were identified, at which point all identified chains were classified into a single cluster. The process was repeated for unclustered chains until all chains were assigned to clusters.

#### Time correlation analysis

The relaxation times of various collective variables were determined by first calculating the autocorrelation function of each variable and then fitting the resulting relaxation curve to an exponential function. For collective variables, we analyzed patch-to-patch distances across the single chain, slab coexistence, and cubic box simulations. In condensate simulations, we reported the average relaxation time over 100 chains. In the cubic box simulations, we additionally examined the coordination number of charged patches, defined as the count of negatively charged patches within a 0.7 nm cutoff distance from any given positive patch. The reported values represent the average relaxation time across all 100 positive patches in the simulation.

### 2. Supporting Figures

**Figure S1.**
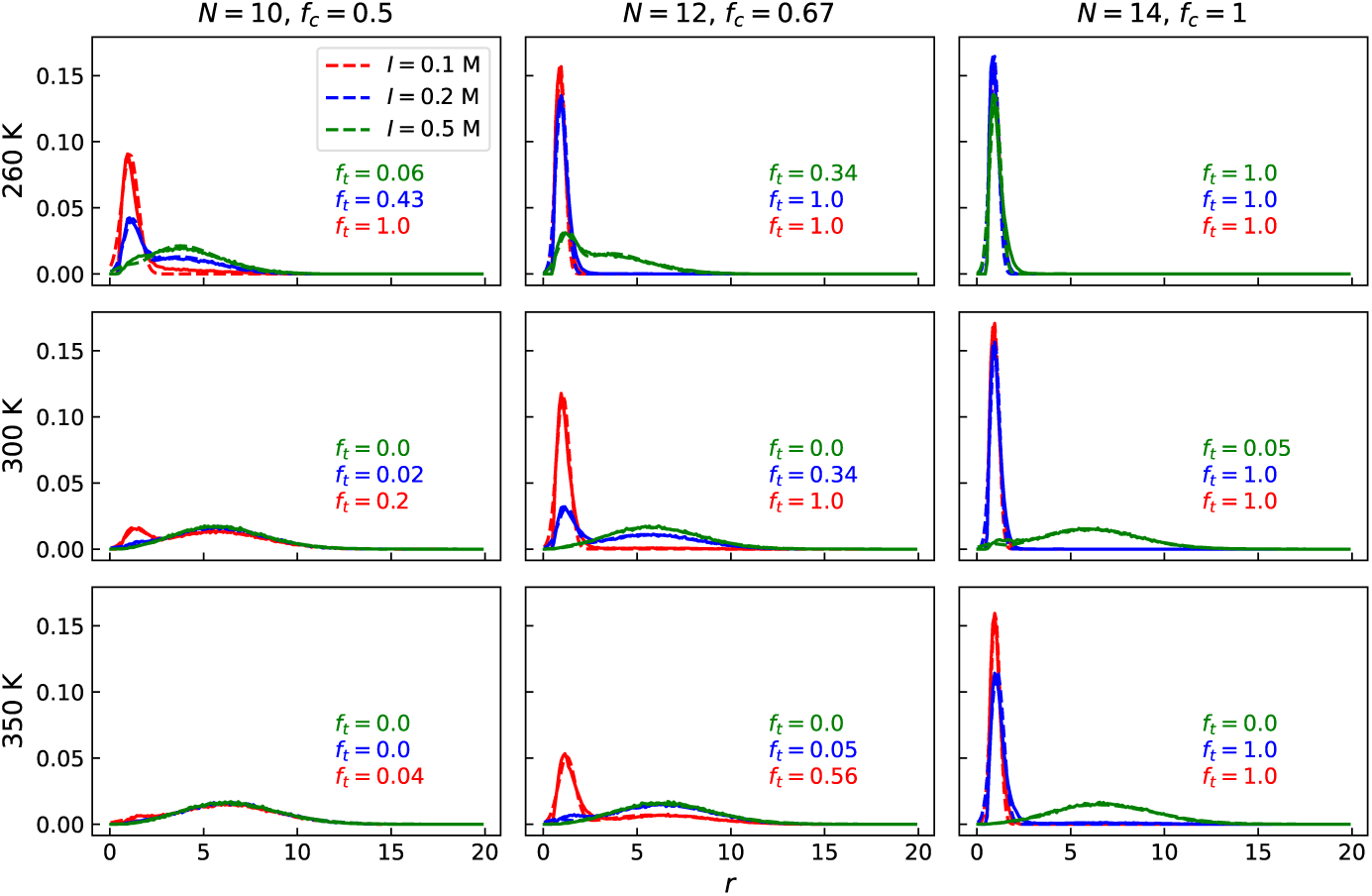
Fittings for determining *f_t_* from simulated patch-to-patch distance distribution *p*(*r*). Solid lines indicate results from simulations while dashed lines show the optimal fitting according to the adjusted polymer model in Eq. 1 of the main text.

**Figure S2.**
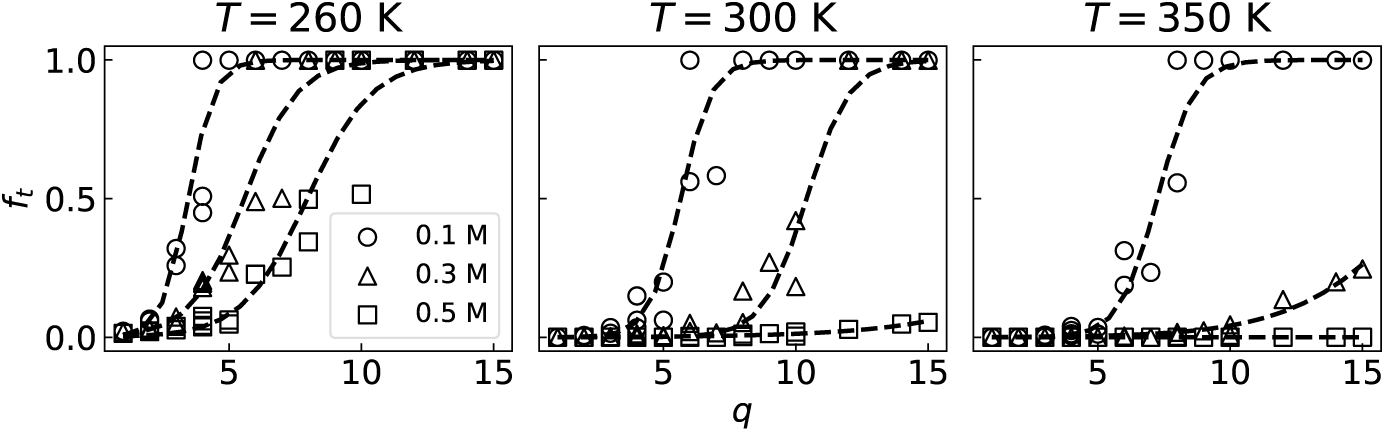
Correlation between total number of charged amino acids within each charge patch *q* = *f_c_N* and *f_t_* for different temperature and ionic strength conditions. Markers indicate results from simulation and dashed lines are the best fit to the sigmoid equation shown in Eq. 4 of the main text.

**Figure S3.**
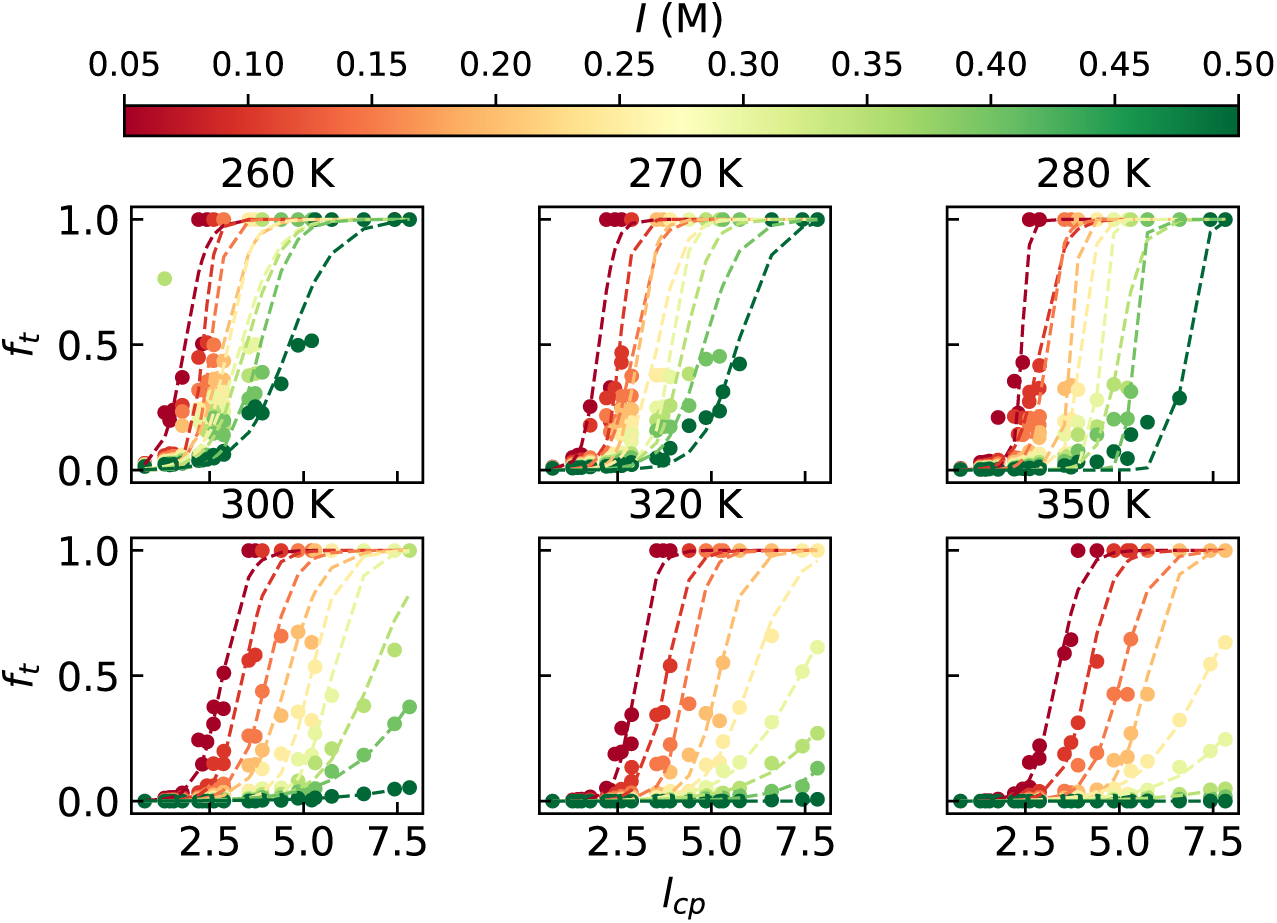
Global fitting of *f_t_* as a function of *l_cp_* for all simulations to obtain an optimized *β* of 0.76. Dots mark the results from simulated distance distribution while dashed lines indicate fitting from a sigmoid function shown in Eq. 6 of the main text.

**Figure S4.**
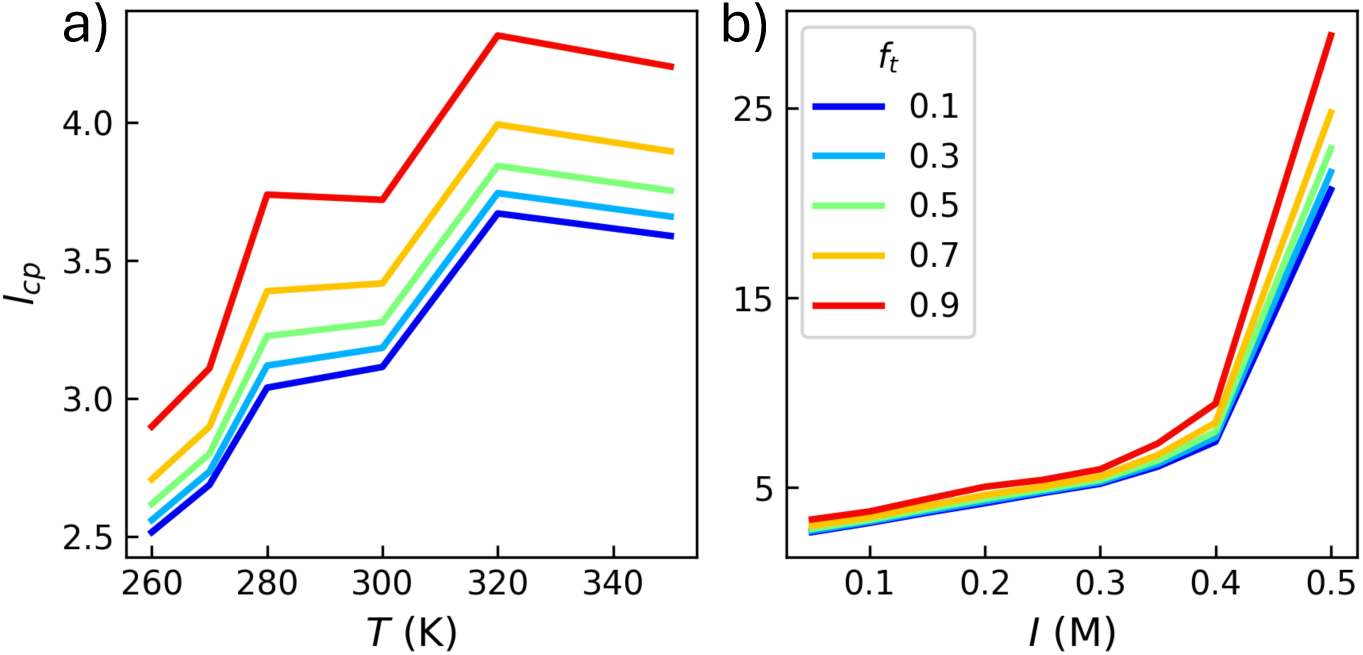
The value of *l_cp_* for which *f_t_* achieves a given value at a) different temperatures and a fixed ionic strength of 0.1M and at b) different ionic strengths and a fixed temperature of 300K.

**Figure S5.**
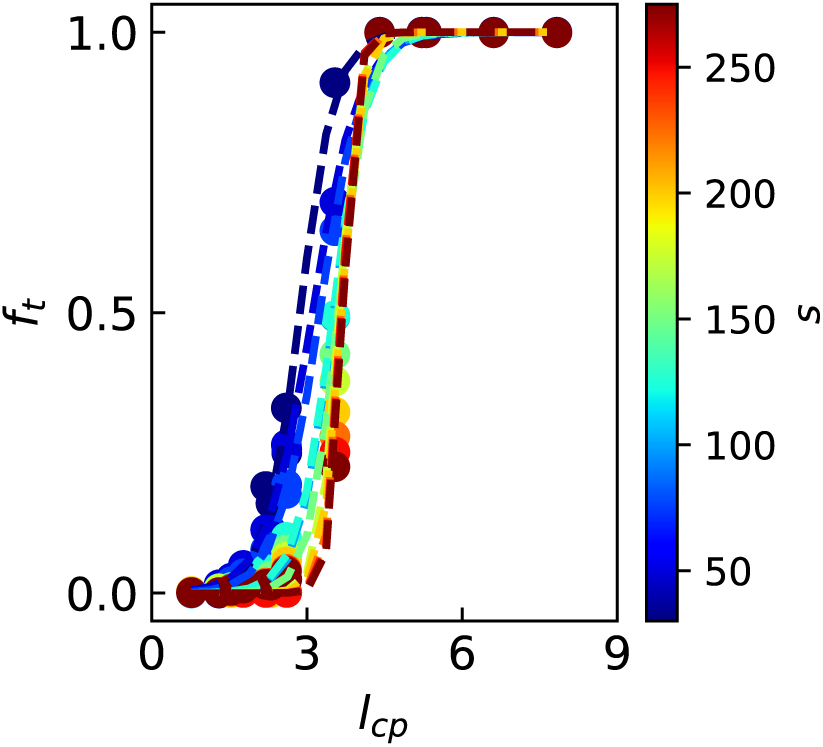
*f_t_* as a function of *l_cp_* for different patch sequence separations *s*. Filled dots show results from simulation while Dashed lines indicate the best fit results from the sigmoid function shown in Eq. 6 of the main text.

**Figure S6.**
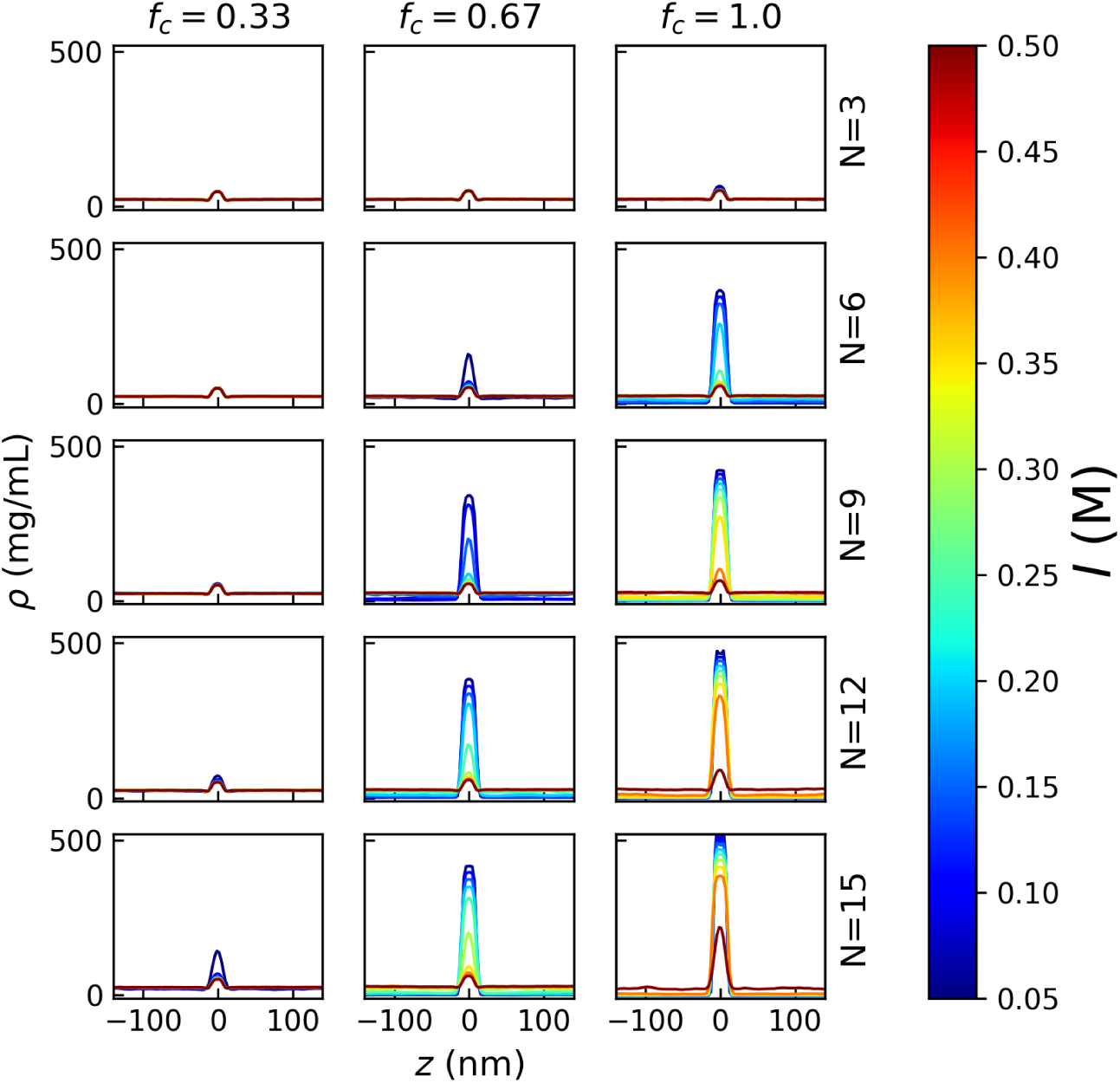
Slab coexistence density profiles as a function of the charge content (*N* and *f_c_*) and the ionic strength *I*.

**Figure S7.**
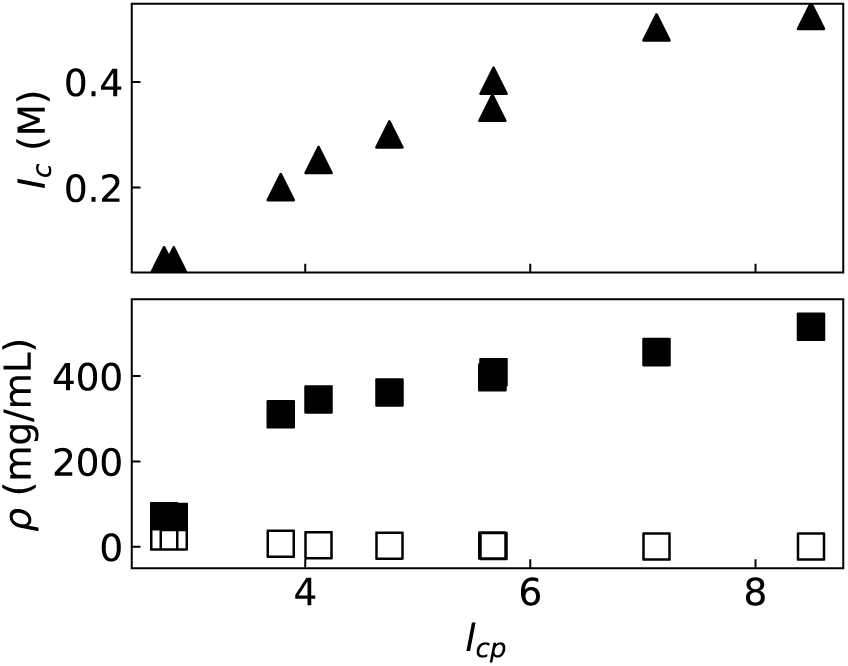
Critical ionic strength *I_c_* as a function of *l_cp_*. Dense phase (filled) and saturation concentrations (empty) as a function of *l_cp_*.

**Figure S8.**
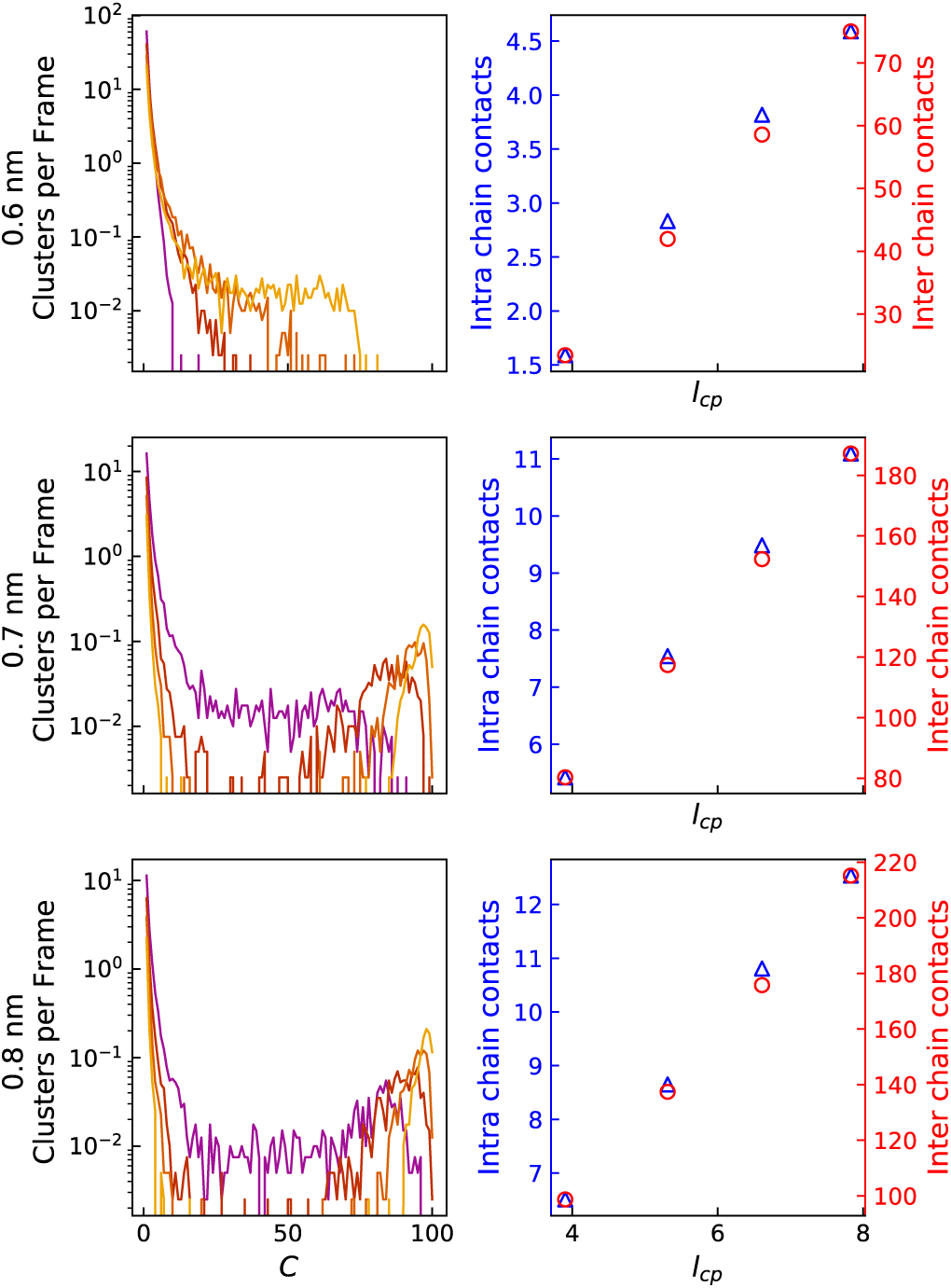
Clustering analysis (left) and number of intra (blue) and inter (red) chain charge contacts (right) for the cubic box simulations at different *l_cp_* following the same color code as Fig. 3 in the main text. Chains are only considered in contact when at least one pair of residues from the positive and negative charged segments have a distance less than the displayed cutoff distance.

**Figure S9.**
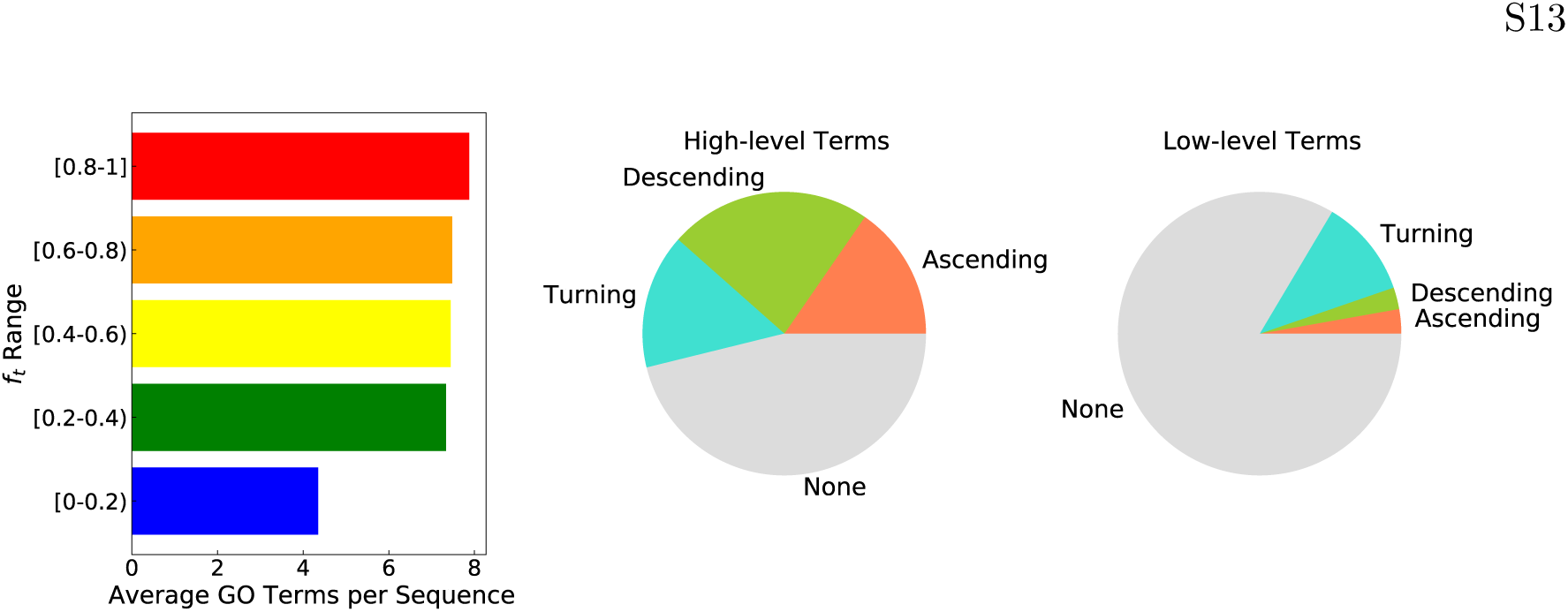
**a)** Number of annotated GO terms per sequence in each *f_t_* bin from regions of the IDRome database with segregated positive and negative charges. **b)** Relative population of GO terms in each dependence class for high level and low level terms.

### 3. Supporting Tables

**Table S1.** The amino acid parameters used in the model. *σ* is the diameter of the amino acid used in the short-ranged pair potential. *λ* is the scaled hydropathy from the literature. [2]

| Type | Mass (amu) | Charge | $\sigma$ (Å) | $\lambda$ |
| --- | --- | --- | --- | --- |
| ARG | 156.20 | 1 | 6.56 | 0.000 |
| ASP | 115.10 | -1 | 5.58 | 0.378 |
| GLY | 57.05 | 0 | 4.50 | 0.649 |

**Table S2.** List of sequence charge content (*N* and *f_c_*), temperature (*T*), and ionic strength (*I*) for the single-chain simulations at a fixed sequence separation *s*=100.

| Single chain simulations (s=100 residues) |  |  |  |  |  |  |  |  |  |
| --- | --- | --- | --- | --- | --- | --- | --- | --- | --- |
| $N$ | 3 | 4 | 6 | 8 | 9 | 10 | 12 | 14 | 15 |
| $f_c$ | 1/3,<br>2/3, 1 | 1/2, 1 | 1/3,<br>2/3, 1 | 1/2, 1 | 1/3,<br>2/3, 1 | 1/2, 1 | 1/3,<br>2/3, 1 | 1/2, 1 | 1/3,<br>2/3, 1 |
| $T$ (K) | 260, 270, 280, 300, 320, 350 | | | | | | | | |
| $I$ (M) | 0.05, 0.1, 0.15, 0.2, 0.25, 0.3, 0.35, 0.4, 0.5 | | | | | | | | |

**Table S3.** List of sequence charge content (*N* and *f_c_*), temperature (*T*), and ionic strength (*I*) for the single-chain simulations at different sequence separations.

| Sequence separation simulations |  |  |  |  |  |
| --- | --- | --- | --- | --- | --- |
| $N$ | 3 | 6 | 9 | 12 | 15 |
| $f_c$ | 1/3, 2/3, 1 | 1/3, 2/3, 1 | 1/3, 2/3, 1 | 1/3, 2/3, 1 | 1/3, 2/3, 1 |
| $s$ | 30, 50, 75, 100, 125, 150, 175, 200, 225, 250, 275, 300 | | | | |
| $T$ (K) | 300 | | | | |
| $I$ (M) | 0.1 | | | | |

**Table S4.** List of sequence charge content (*N* and *f_c_*), temperature (*T*), and ionic strength (*I*) for the coexistence slab simulations.

|  |  |  |  |  |  |  |  |  |  |
| --- | --- | --- | --- | --- | --- | --- | --- | --- | --- |
| $N$ | 3 | 4 | 6 | 8 | 9 | 10 | 12 | 14 | 15 |
| $f_c$ | 1/3,<br>2/3, 1 | 1/2, 1 | 1/3,<br>2/3, 1 | 1/2, 1 | 1/3,<br>2/3, 1 | 1/2, 1 | 1/3,<br>2/3, 1 | 1/2, 1 | 1/3,<br>2/3, 1 |
| $T$ (K) | 280 | | | | | | | | |
| $I$ (M) | 0.05, 0.1, 0.15, 0.2, 0.25, 0.3, 0.35, 0.4, 0.5 | | | | | | | | |

**Table S5.** List of sequence charge content (*N* and *f_c_*), temperature (*T*), and ionic strength (*I*) for the cubic box simulations.

| **Cubic box simulations** |  |  |  |  |
| --- | --- | --- | --- | --- |
| $N$ | 6 | 9 | 12 | 15 |
| $f_c$ | 1 | 2/3, 1 | 2/3, 1 | 2/3, 1 |
| $T$ (K) | 280 | | | |
| $I$ (M) | 0.1 | | | |

## References

(1) Tompa, P. Intrinsically disordered proteins: a 10-year recap. Trends Biochem. Sci. 2012, 37, 509–516.

(2) Uversky, V. N. The multifaceted roles of intrinsic disorder in protein complexes. FEBS letters 2015, 589, 2498–2506.

(3) Wright, P. E.; Dyson, H. J. Intrinsically disordered proteins in cellular signalling and regulation. Nat. Rev. Mol. Cell Biol. 2015, 16, 18–29.

(4) Elbaum-Garfinkle, S.; Kim, Y.; Szczepaniak, K.; Chen, C. C.-H.; Eckmann, C. R.; Myong, S.; Brangwynne, C. P. The disordered P granule protein LAF-1 drives phase separation into droplets with tunable viscosity and dynamics. Proc. Natl. Acad. Sci. U.S.A. 2015, 112, 7189–7194.

(5) Burke, K. A.; Janke, A. M.; Rhine, C. L.; Fawzi, N. L. Residue-by-residue view of in vitro FUS granules that bind the C-terminal domain of RNA polymerase II. Mol. Cell 2015, 60, 231–241.

(6) Alberti, S.; Gladfelter, A.; Mittag, T. Considerations and challenges in studying liquid-liquid phase separation and biomolecular condensates. Cell 2019, 176, 419–434.

(7) Avni, A.; Swasthi, H. M.; Majumdar, A.; Mukhopadhyay, S. Intrinsically disordered proteins in the formation of functional amyloids from bacteria to humans. Progress in Molecular Biology and Translational Science 2019, 166, 109–143.

(8) Louros, N.; Schymkowitz, J.; Rousseau, F. Mechanisms and pathology of protein misfolding and aggregation. Nature Reviews Molecular Cell Biology 2023, 24, 912–933.

(9) Eliezer, D. Biophysical characterization of intrinsically disordered proteins. Curr. Opin. Struct. Biol. 2009, 19, 23–30.

(10) Best, R. B. Computational and theoretical advances in studies of intrinsically disordered proteins. Curr. Opin. Struct. Biol. 2017, 42, 147–154.

(11) Onuchic, J. N.; Wolynes, P. G. Theory of protein folding. Curr. Opin. Struct. Biol. 2004, 14, 70–75.

(12) Dill, K. A.; MacCallum, J. L. The protein-folding problem, 50 years on. Science 2012, 338, 1042–1046.

(13) Abramson, J.; Adler, J.; Dunger, J.; Evans, R.; Green, T.; Pritzel, A.; Ronneberger, O.; Willmore, L.; Ballard, A. J.; Bambrick, J., et al. Accurate structure prediction of biomolecular interactions with AlphaFold 3. Nature 2024, 630, 493–500.

(14) Borgia, A.; Zheng, W.; Buholzer, K.; Borgia, M. B.; Schuler, A.; Hofmann, H.; Soranno, A.; Nettels, D.; Gast, K.; Grishaev, A.; Best, R. B.; Schuler, B. Consistent view of polypeptide chain expansion in chemical denaturants from multiple experimental methods. J. Am. Chem. Soc. 2016, 138, 11714–11726.

(15) Brookes, D. H.; Head-Gordon, T. Experimental inferential structure determination of ensembles for intrinsically disordered proteins. J. Am. Chem. Soc. 2016, 138, 4530–4538.

(16) Köfinger, J.; Stelzl, L. S.; Reuter, K.; Allande, C.; Reichel, K.; Hummer, G. Efficient ensemble refinement by reweighting. J. Chem. Theory Comput. 2019, 15, 3390–3401.

(17) Lazar, T.; Martínez-Pérez, E.; Quaglia, F.; Hatos, A.; Chemes, L. B.; Iserte, J. A.; Méndez, N. A.; Garrone, N. A.; Saldaño, T. E.; Marchetti, J., et al. PED in 2021: a major update of the protein ensemble database for intrinsically disordered proteins. Nucleic Acids Res. 2021, 49, D404–D411.

(18) Gomes, G.-N. W.; Namini, A.; Gradinaru, C. C. Integrative conformational ensembles of Sic1 using different initial pools and optimization methods. Front. Mol. Biosci. 2022, 9, 910956.

(19) Das, R. K.; Pappu, R. V. Conformations of intrinsically disordered proteins are influenced by linear sequence distributions of oppositely charged residues. Proc. Natl. Acad. Sci. U.S.A. 2013, 110, 13392–13397.

(20) Sawle, L.; Ghosh, K. A theoretical method to compute sequence dependent configurational properties in charged polymers and proteins. J. Chem. Phys. 2015, 143, 085101.

(21) Zheng, W.; Dignon, G.; Brown, M.; Kim, Y. C.; Mittal, J. Hydropathy patterning complements charge patterning to describe conformational preferences of disordered proteins. J. Phys. Chem. Lett. 2020, 11, 3408–3415.

(22) Holehouse, A. S.; Kragelund, B. B. The molecular basis for cellular function of intrinsically disordered protein regions. Nat. Rev. Mol. Cell Biol. 2024, 25, 187–211.

(23) Tesei, G.; Trolle, A. I.; Jonsson, N.; Betz, J.; Knudsen, F. E.; Pesce, F.; Johansson, K. E.; Lindorff-Larsen, K. Conformational ensembles of the human intrinsically disordered proteome. Nature 2024, 626, 897–904.

(24) Saar, K. L.; Morgunov, A. S.; Qi, R.; Arter, W. E.; Krainer, G.; Lee, A. A.; Knowles, T. P. Learning the molecular grammar of protein condensates from sequence determinants and embeddings. Proceedings of the National Academy of Sciences 2021, 118, e2019053118.

(25) Bremer, A.; Farag, M.; Borcherds, W. M.; Peran, I.; Martin, E. W.; Pappu, R. V.; Mittag, T. Deciphering how naturally occurring sequence features impact the phase behaviours of disordered prion-like domains. Nature Chemistry 2022, 14, 196–207.

(26) Alshareedah, I.; Borcherds, W. M.; Cohen, S. R.; Singh, A.; Posey, A. E.; Farag, M.; Bremer, A.; Strout, G. W.; Tomares, D. T.; Pappu, R. V., et al. Sequence-specific interactions determine viscoelasticity and ageing dynamics of protein condensates. Nat. Phys. 2024, 20, 1482–1491.

(27) Riback, J. A.; Katanski, C. D.; Kear-Scott, J. L.; Pilipenko, E. V.; Rojek, A. E.; Sosnick, T. R.; Drummond, D. A. Stress-triggered phase separation is an adaptive, evolutionarily tuned response. Cell 2017, 168, 1028–1040.

(28) Du, Z.; Wang, H.; Wu, C.; Buck, M.; Zheng, W.; Hansen, A. L.; Kao, H.-Y.; Yang, S. Phosphorylation modulates estrogen receptor disorder by altering long-range hydrophobic interactions. https://www.biorxiv.org/content/10.1101/2023.07.14.548966v1 2023,

(29) Maiti, S.; Maji, T.; Saibo, N. V.; De, S., et al. Experimental methods to study the structure and dynamics of intrinsically disordered regions in proteins. Curr. Opin. Struct. Biol. 2024, 100138.

(30) Kikhney, A. G.; Svergun, D. I. A practical guide to small angle X-ray scattering (SAXS) of flexible and intrinsically disordered proteins. FEBS Lett. 2015, 589, 2570–2577.

(31) Gast, K.; Fiedler, C. Dynamic and static light scattering of intrinsically disordered proteins. Intrinsically Disordered Protein Analysis: Volume 2, Methods and Experimental Tools 2012, 137–161.

(32) Choy, W.-Y.; Mulder, F. A. A.; Crowhurst, K. A.; Muhandiram, D. R.; Millet, I. S.; Doniach, S.; Forman-Kay, J. D.; Kay, L. E. Distribution of molecular size within an unfolded state ensemble using small-angle X-ray scattering and pulse field gradient NMR techniques. J. Mol. Biol. 2002, 316, 101–112.

(33) Schuler, B.; Soranno, A.; Hofmann, H.; Nettels, D. Single-molecule FRET spectroscopy and the polymer physics of unfolded and intrinsically disordered proteins. Annu. Rev. Biophys. 2016, 45, 207–231.

(34) Kjaergaard, M.; Norholm, A.-B.; Hendus-Altenburger, R.; Pedersen, S. F.; Poulsen, F. M.; Kragelund, B. B. Temperature-dependent structural changes in intrinsically-disordered proteins: formation of *α*-helices or loss of polyproline II? Protein Sci. 2010, 19, 1555–1564.

(35) Wuttke, R.; Hofmann, H.; Nettels, D.; Borgia, M. B.; Mittal, J.; Best, R. B.; Schuler, B. Temperature-dependent solvation modulates the dimensions of disordered proteins. Proc. Natl. Acad. Sci. U.S.A. 2014, 111, 5213–5218.

(36) Wiggers, F.; Wohl, S.; Dubovetskyi, A.; Rosenblum, G.; Zheng, W.; Hofmann, H. Diffusion of a disordered protein on its folded ligand. Proc. Natl. Acad. Sci. U.S.A. 2021, 118, e2106690118.

(37) König, I.; Soranno, A.; Nettels, D.; Schuler, B. Impact of in-cell and in-vitro crowding on the conformations and dynamics of an intrinsically disordered protein. Angew. Chem. 2021, 133, 10819–10824.

(38) Dignon, G. L.; Zheng, W.; Kim, Y. C.; Best, R. B.; Mittal, J. Sequence determinants of protein phase behavior from a coarse-grained model. PLoS Comput. Biol. 2018, 14, e1005941.

(39) Joseph, J. A.; Reinhardt, A.; Aguirre, A.; Chew, P. Y.; Russell, K. O.; Espinosa, J. R.; Garaizar, A.; Collepardo-Guevara, R. Physics-driven coarse-grained model for biomolecular phase separation with near-quantitative accuracy. Nature Computational Science 2021, 1, 732–743.

(40) Marsh, J. A.; Singh, V. K.; Jia, Z.; Forman-Kay, J. D. Sensitivity of secondary structure propensities to sequence differences between *α*-and *γ*-synuclein: implications for fibrillation. Protein Science 2006, 15, 2795–2804.

(41) Englander, S. W. Protein folding intermediates and pathways studied by hydrogen exchange. Annu. Rev. Biophys. Biomol. Struct. 2000, 29, 213–238.

(42) Xu, G.; Chance, M. R. Hydroxyl radical-mediated modification of proteins as probes for structural proteomics. Chemical reviews 2007, 107, 3514–3543.

(43) Hartlmüller, C.; Spreitzer, E.; Göbl, C.; Falsone, F.; Madl, T. NMR characterization of solvent accessibility and transient structure in intrinsically disordered proteins. J. Biomol. NMR 2019, 73, 305–317.

(44) Hansen, D. F.; Feng, H.; Zhou, Z.; Bai, Y.; Kay, L. E. Selective characterization of microsecond motions in proteins by NMR relaxation. J. Am. Chem. Soc. 2009, 131, 16257–16265.

(45) Kjaergaard, M.; Poulsen, F. M. Disordered proteins studied by chemical shifts. Prog. Nucl. Magn. Reson. Spectrosc. 2012, 60, 42–51.

(46) Clore, G. M. Practical aspects of paramagnetic relaxation enhancement in biological macromolecules. Methods Enzymol. 2015, 564, 485–497.

(47) Schuler, B.; Hofmann, H. Single-molecule spectroscopy of protein folding dynamics—expanding scope and timescales. Curr. Opin. Struct. Biol. 2013, 23, 36–47.

(48) Lapidus, L. J.; Eaton, W. A.; Hofrichter, J. Measuring the rate of intramolecular contact formation in polypeptides. Proc. Natl. Acad. Sci. U.S.A. 2000, 97, 7220–7225.

(49) Marsh, J. A.; Forman-Kay, J. D. Sequence determinants of compaction in intrinsically disordered proteins. Biophys. J. 2010, 98, 2383–2390.

(50) Flory, P. J. Principles of polymer chemistry; Cornell university press, 1953.

(51) de Gennes, P.-G. Scaling Concepts in Polymer Physics; Cornell University Press, 1978.

(52) Zheng, W.; Zerze, G. H.; Borgia, A.; Mittal, J.; Schuler, B.; Best, R. B. Inferring properties of disordered chains from FRET transfer efficiencies. J. Chem. Phys. 2018, 148, 123329.

(53) Zheng, W.; Best, R. B. An extended Guinier analysis for intrinsically disordered proteins. J. Mol. Biol. 2018, 430, 2540–2553.

(54) Dignon, G. L.; Zheng, W.; Best, R. B.; Kim, Y. C.; Mittal, J. Relation between single-molecule properties and phase behavior of intrinsically disordered proteins. Proc. Natl. Acad. Sci. U.S.A. 2018, 115, 9929–9934.

(55) Martin, E. W.; Holehouse, A. S.; Peran, I.; Farag, M.; Incicco, J. J.; Bremer, A.; Grace, C. R.; Soranno, A.; Pappu, R. V.; Mittag, T. Valence and patterning of aromatic residues determine the phase behavior of prion-like domains. Science 2020, 367, 694–699.

(56) Salmon, L.; Nodet, G.; Ozenne, V.; Yin, G.; Jensen, M. R.; Zweckstetter, M.; Blackledge, M. NMR characterization of long-range order in intrinsically disordered proteins. J. Am. Chem. Soc. 2010, 132, 8407–8418.

(57) Otteson, L.; Nagy, G.; Kunkel, J.; Kodis, G.; Zheng, W.; Bignon, C.; Longhi, S.; Grubmüller, H.; Vaiana, A. C.; Vaiana, S. M. Transient Non-local Interactions Dominate the Dynamics of Measles Virus NTAIL. https://www.biorxiv.org/content/10.1101/2024.07.22.604679v1 2024,

(58) Peng, Y. et al. A metastable contact and structural disorder in the estrogen receptor transactivation domain. Structure 2019, 27, 229–240.

(59) Zheng, W.; Du, Z.; Ko, S. B.; Wickramasinghe, N. P.; Yang, S. Incorporation of D2O-Induced Fluorine Chemical Shift Perturbations into Ensemble-Structure Characterization of the ERalpha Disordered Region. J. Phys. Chem. B 2022, 126, 9176–9186.

(60) Saibo, N. V.; Maiti, S.; Boral, S.; Banerjee, P.; Kushwaha, T.; Inampudi, K. K.; Goswami, R.; De, S. The intrinsically disordered transactivation region of HOXA9 regulates its function by auto-inhibition of its DNA-binding activity. International Journal of Biological Macromolecules 2024, 273, 132704.

(61) Martinez-Yamout, M. A.; Nasir, I.; Shnitkind, S.; Ellis, J. P.; Berlow, R. B.; Kroon, G.; Deniz, A. A.; Dyson, H. J.; Wright, P. E. Glutamine-rich regions of the disordered CREB transactivation domain mediate dynamic intra-and intermolecular interactions. Proceedings of the National Academy of Sciences 2023, 120, e2313835120.

(62) Mohanty, P.; Phan, T. M.; Mittal, J. Transient interdomain interactions modulate the monomeric structural ensemble and oligomerization landscape of Huntingtin Exon 1. https://www.biorxiv.org/content/10.1101/2024.05.03.592468v1 2024,

(63) Mohanty, P.; Rizuan, A.; Kim, Y. C.; Fawzi, N. L.; Mittal, J. A complex network of interdomain interactions underlies the conformational ensemble of monomeric TDP-43 and modulates its phase behavior. Protein Science 2024, 33, e4891.

(64) Krois, A. S.; Dyson, H. J.; Wright, P. E. Long-range regulation of p53 DNA binding by its intrinsically disordered N-terminal transactivation domain. Proc. Natl. Acad. Sci. U.S.A. 2018, 115, E11302–E11310.

(65) Choi, J.-M.; Hyman, A. A.; Pappu, R. V. Generalized models for bond percolation transitions of associative polymers. *Phys*. Rev. E 2020, 102, 042403.

(66) Das, S.; Muthukumar, M. Microstructural Organization in *α*-Synuclein Solutions. Macromolecules 2022, 55, 4228–4236.

(67) Zheng, W.; Borgia, A.; Buholzer, K.; Grishaev, A.; Schuler, B.; Best, R. B. Probing the action of chemical denaturant on an intrinsically disordered protein by simulation and experiment. J. Am. Chem. Soc. 2016, 138, 11702–11713.

(68) Zeng, X.; Ruff, K. M.; Pappu, R. V. Competing interactions give rise to two-state behavior and switch-like transitions in charge-rich intrinsically disordered proteins. Proc. Natl. Acad. Sci. U.S.A. 2022, 119, e2200559119.

(69) Wohl, S.; Zheng, W. Interpreting Transient Interactions of Intrinsically Disordered Proteins. J. Phys. Chem. B 2023, 127, 2395–2406.

(70) Des Cloizeaux, J. Lagrangian theory for a self-avoiding random chain. Phys. Rev. A 1974, 10, 1665.

(71) Le Guillou, J.; Zinn-Justin, J. Critical exponents for the n-vector model in three dimensions from field theory. Phys. Rev. Lett. 1977, 39, 95.

(72) Fisher, M. E. Shape of a Self-Avoiding Walk or Polymer Chain. J. Chem. Phys. 1966, 44, 616–622.

(73) Hofmann, H.; Soranno, A.; Borgia, A.; Gast, K.; Nettels, D.; Schuler, B. Polymer scaling laws of unfolded and intrinsically disordered proteins quantified with single-molecule spectroscopy. Proc. Natl. Acad. Sci. U.S.A. 2012, 109, 16155–16160.

(74) Huihui, J.; Firman, T.; Ghosh, K. Modulating charge patterning and ionic strength as a strategy to induce conformational changes in intrinsically disordered proteins. J. Chem. Phys. 2018, 149, 085101.

(75) Amin, A. N.; Lin, Y.-H.; Das, S.; Chan, H. S. Analytical theory for sequence-specific binary fuzzy complexes of charged intrinsically disordered proteins. J. Phys. Chem. B 2020, 124, 6709–6720.

(76) von Bülow, S.; Tesei, G.; Lindorff-Larsen, K. Prediction of phase separation propensities of disordered proteins from sequence. https://www.biorxiv.org/content/10.1101/2024.06.03.597109v1 2024,

(77) Mittag, T.; Pappu, R. V. A conceptual framework for understanding phase separation and addressing open questions and challenges. Mol. Cell 2022, 82, 2201–2214.

(78) Sternke-Hoffmann, R.; Sun, X.; Dos Santos Pinto, M.; Venclovaite, U.; Menzel, A.; Wördehoff, M.; Hoyer, W.; Zheng, W.; Luo, J. Phase separation and aggregation of *α*-synuclein diverge at different salt conditions. Adv. Sci. 2024, 11, 2308279.

(79) Wang, J.; Devarajan, D. S.; Kim, Y. C.; Nikoubashman, A.; Mittal, J. Sequence-Dependent Conformational Transitions of Disordered Proteins During Condensation. https://www.biorxiv.org/content/10.1101/2024.01.11.575294v1 2024,

(80) Joshi, A.; Walimbe, A.; Avni, A.; Rai, S. K.; Arora, L.; Sarkar, S.; Mukhopadhyay, S. Single-molecule FRET unmasks structural subpopulations and crucial molecular events during FUS low-complexity domain phase separation. Nature Communications 2023, 14, 7331.

(81) Pal, T.; Wesśen, J.; Das, S.; Chan, H. S. Differential Effects of Sequence-Local versus Nonlocal Charge Patterns on Phase Separation and Conformational Dimensions of Polyampholytes as Model Intrinsically Disordered Proteins. J. Phys. Chem. Lett. 2024, 15, 8248–8256.

(82) Consortium, G. O. The Gene Ontology (GO) database and informatics resource. Nucleic acids research 2004, 32, D258–D261.

(83) Ruggeri, F.; Zosel, F.; Mutter, N.; Różycka, M.; Wojtas, M.; Ożyhar, A.; Schuler, B.; Krishnan, M. Single-molecule electrometry. Nat. Nanotechnol. 2017, 12, 488–495.

(84) Krishnan, M. A simple model for electrical charge in globular macromolecules and linear polyelectrolytes in solution. J. Chem. Phys. 2017, 146.

(85) Aso, T.; Vasavada, H. A.; Kawaguchi, T.; Germino, F. J.; Ganguly, S.; Kitajima, S.; Weissman, S. M.; Yasukochi, Y. Characterization of cDNA for the large subunit of the transcription initiation factor TFIIF. Nature 1992, 355, 461–464.

## References

[1] Dignon, G. L.; Zheng, W.; Kim, Y. C.; Best, R. B.; Mittal, J. PLoS Comput. Biol. 2018, 14, e1005941.

[2] Kapcha, L. H.; Rossky, P. J. J. Mol. Biol. 2014, 426, 484–498.

[3] Debye, P.; Hückel, E. Physikalische Zeitschrift 1923, 24, 185–206.

[4] Ashbaugh, H. S.; Hatch, H. W. J. Am. Chem. Soc. 2008, 130, 9536–9542.

[5] Anderson, J. A.; Glaser, J.; Glotzer, S. C. Comput. Mater. Sci. 2020, 173, 109363.

[6] azplugins; https://github.com/mphowardlab/azplugins; 2024.

